# Functional enhancer elements drive subclass-selective expression from mouse to primate neocortex

**DOI:** 10.1101/555318

**Authors:** John K. Mich, Lucas T. Graybuck, Erik E. Hess, Joseph T. Mahoney, Yoshiko Kojima, Yi Ding, Saroja Somasundaram, Jeremy A. Miller, Natalie Weed, Victoria Omstead, Yemeserach Bishaw, Nadiya V. Shapovalova, Refugio A. Martinez, Olivia Fong, Shenqin Yao, Marty Mortrud, Peter Chong, Luke Loftus, Darren Bertagnolli, Jeff Goldy, Tamara Casper, Nick Dee, Ximena Opitz-Araya, Ali Cetin, Kimberly A. Smith, Ryder P. Gwinn, Charles Cobbs, Andrew. L. Ko, Jeffrey G. Ojemann, C. Dirk Keene, Daniel. L. Silbergeld, Susan M. Sunkin, Viviana Gradinaru, Gregory D. Horwitz, Hongkui Zeng, Bosiljka Tasic, Ed S. Lein, Jonathan T. Ting, Boaz P. Levi

## Abstract

Viral genetic tools to target specific brain cell types in humans and non-genetic model organisms will transform basic neuroscience and targeted gene therapy. Here we used comparative epigenetics to identify thousands of human neuronal subclass-specific putative enhancers to regulate viral tools, and 34% of these were conserved in mouse. We established an AAV platform to evaluate cellular specificity of functional enhancers by multiplexed fluorescent in situ hybridization (FISH) and single cell RNA sequencing. Initial testing in mouse neocortex yields a functional enhancer discovery success rate of over 30%. We identify enhancers with specificity for excitatory and inhibitory classes and subclasses including PVALB, LAMP5, and VIP/LAMP5 cells, some of which maintain specificity *in vivo* or *ex vivo* in monkey and human neocortex. Finally, functional enhancers can be proximal or distal to cellular marker genes, conserved or divergent across species, and could yield brain-wide specificity greater than the most selective marker genes.

## Introduction

A major goal in neuroscience is to establish the distinct role of each cell type in brain circuitry, how they give rise to complex function, and how their dysfunction can cause disease. To accomplish this goal, more precise and widely useful genetic tools for specific experimental study of these cell types across organisms are needed.

Mouse and human neocortex have a remarkable diversity of cell types (Zeisel et al., 2015; Tasic et al., 2016, 2018; Boldog et al., 2018; Hodge et al., 2019, 2020; Bakken et al., 2020). Comparison of gene expression data between mouse and human shows strong conservation of molecular features across major classes (eg. inhibitory, excitatory and glial classes) and subclasses (e.g. medial ganglionic eminence [MGE]-derived PVALB and SST, and caudal ganglionic eminence [CGE]-derived VIP and LAMP5 inhibitory subclasses, layer 2/3 [L2/3] and L4 excitatory subclasses, Hodge et al., 2019). However, many more differences are seen at the level of cell types, and one cell type in human often cannot be unambiguously assigned to an orthologous cell type in mouse (eg. Inh L5-6 *PVALB LGR5*, Hodge et al., 2019). These molecular differences possibly contribute to clinical failures when translating therapeutic approaches from mouse to human (Doody et al., 2014; Egan et al., 2018).

Both similarities and differences in gene expression are regulated by the function of enhancers and other epigenetic elements. Epigenetic profiling with single-cell resolution techniques matched across multiple organisms will allow direct comparison of the regulatory landscapes of cell classes, subclasses and types. Although detailed single cell Assay for Transposase Accessible Chromatin with sequencing (scATAC-seq) datasets profiling mouse brain now exist (Preissl et al., 2018; Cusanovich et al., 2018; Fang et al., 2019; Lareau et al., 2019), few human-specific epigenetic datasets with single cell resolution exist (Luo et al., 2017; Fullard et al., 2018; Lake et al., 2018), and no high-resolution single nucleus (sn) ATAC-seq datasets were available from human neocortex that could be compared directly to mouse. More high-quality human snATAC-seq data (Bakken et al., 2020) will reveal the regulatory elements that confer cell type molecular identity and will enable their specific genetic access via viral vectors.

Genetic tools are vital to establishing the function of cell populations in model organisms, and for testing the function of specific cell classes, subclasses and types. Germline transgenic mice with recombinase-based drivers and reporters allow precision in cell profiling and functional experimentation not possible in non-genetically tractable organisms. Recent studies have tried to overcome this barrier using viral vectors for transgene delivery (Dimidschstein et al., 2016; Ting et al., 2018). However, few tools are available to allow targeted gene expression at cell subclass or type resolution—particularly in primates, a model system with unique advantages for neuroscience. A new toolkit is required to access and interrogate those subclasses and types.

Adeno-associated viruses (AAVs) are ubiquitous and non-pathogenic viruses that allow transduction of adult post-mitotic neurons and could be leveraged to build tools for genetic access to specific brain cell subclasses. AAV capsids have been identified that possess unique tropisms like retrograde transport (Tervo et al., 2016), and blood brain barrier penetrance (Deverman et al., 2016; Chan et al., 2017), and can be exploited to deliver specific transgenes in many tissues (Deverman et al., 2016; Greig et al., 2018; Song et al., 2019). Specific promoters and enhancers can be used to further control transgene expression (Nord et al., 2013; Visel et al., 2013; Silberberg et al., 2016; Dimidschstein et al., 2016; Xiong et al., 2019; Jüttner et al., 2019; Nair et al., 2020). The genome capacity of AAV is ~4.7kb, so only small gene regulatory elements (i.e. enhancers or promoters) can be used for selective targeting of gene expression, and few suitably compact cell class or subclass-specific regulatory elements are known that function across mammalian species (Dimidschstein et al., 2016; Mehta et al., 2019). A set of compact enhancers that confer selective transgene expression in the brains of multiple species including humans will help realize the promise of AAVs for manipulating specific brain cell classes, subclasses, and types.

Here we present a multistep process to generate AAV vectors that drive cell subclass-specific reporter expression across species. First, we conducted snATAC-seq from human temporal cortex. Second, we mined the data to reveal putative subclass-selective enhancers. Third, we tested enhancer function by reporter expression from AAVs in mouse brain, and fourth, we validated specificity in non-human primate (NHP) *in vivo* and *ex vivo,* and human *ex vivo*. Our epigenetic snATAC-seq data generated a high-quality subclass-resolution human neocortex enhancer catalog. Comparison to a similar mouse dataset revealed conserved and divergent subclass-specific putative regulatory elements we leveraged to build reporter-AAV vectors. We generated a collection of subclass-specific AAV vectors that, after systemic delivery, drove neocortical transgene expression patterns predicted by enhancer accessibility profiles. This system for enhancer identification and validation was efficient (over 30% of enhancers tested showed specificity), and generated collections of vectors that labeled subclasses of excitatory and inhibitory cells including PVALB and LAMP5 inhibitory cell subclasses. Several PVALB-specific vectors labeled *Pvalb^+^* subcortical cell populations, demonstrating that the endogenous expression pattern of *Pvalb* could be parcellated by multiple different enhancers. Finally, we identified AAV vectors that maintained specificity in mouse, NHP, and human neocortical tissue: one that labeled all subclasses of inhibitory cells, another that labeled CGE-derived VIP and LAMP5 subclass interneurons, and several that labeled PVALB^+^ cells. These results provide a path forward for identifying enhancers that function in AAV vectors to drive gene expression in cell classes and subclasses across the brain and across species.

## Results

### Epigenetic analysis of human neurons

To find distinguishing neocortical cell subclass-specific enhancers, we generated high-quality chromatin accessibility profiles from multiple middle temporal gyrus (MTG) neurosurgical specimens that were never frozen (bulk, n = 5; single nucleus, n = 14, Fig. S1), using the assay for transposase-accessible chromatin and high-throughput sequencing (ATAC-seq) (Buenrostro et al., 2015; Gray et al., 2017; Graybuck et al., 2019) on bulk populations (Fig. S2) and sorted single nuclei (Fig. S3). We prepared 3,660 single nucleus (sn) ATAC-seq libraries (median of 48,542 uniquely mapped reads per nucleus), and used 2,858 quality-filtered nuclei for clustering and mapping to human snRNA-seq data (Fig. 1A). We excluded nuclei with fewer than 10,000 unique reads, a TSS enrichment score of <4, or <15% of reads overlapping with known DNaseI hypersensitivity peaks isolated from human prefrontal cortex (The ENCODE Project Consortium, 2012). We defined 27 robustly detectable snATAC-seq clusters (Fig. S4) that were mapped by Cicero to the transcriptomic classification at the level of cell subclasses or cell types (Fig. 1B, S5). Several snATAC-seq clusters mapped to the same subclass (Fig. S5). Overall, the cells mapped to all three major classes of brain cells: excitatory, inhibitory, and non-neuronal, which we subdivided into eleven subclasses: excitatory layer 2/3 (L23), L4, L5/6 intra-telencephalic (L56IT), and deep layer non-intratelencephalic neurons (DL); inhibitory LAMP5, VIP, SST, and PVALB neurons; and non-neuronal astrocytes (Astro), microglia (Micro), and oligodendrocytes/OPCs (OligoOPC). In support of the accuracy of mapping to transcriptomic cell subclasses, nuclei microdissected and sorted from superficially neocortical layers usually mapped to superficial cell subclasses (L23, LAMP5, VIP), nuclei microdissected from deep layers mapped to cells found in subgranular neocortical layers (DL, L56IT), and NeuN-negative cells predominantly mapped to non-neuronal cell subclasses (Fig. 1C and Fig. S5G). Importantly, all subclasses contained nuclei contributed by multiple specimens (Fig. 1D).

**Figure 1.**
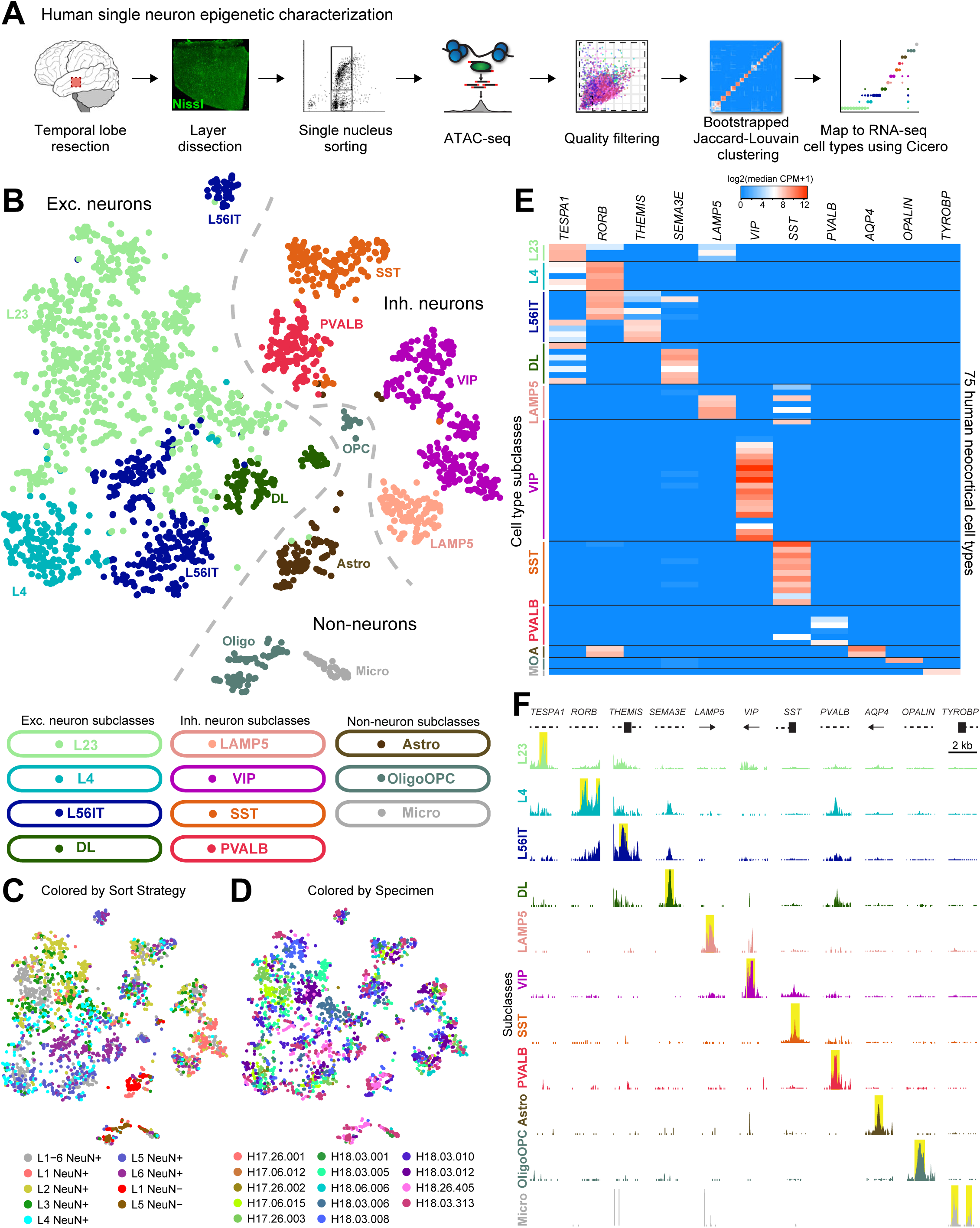
A database of human neocortical cell subclass-specific accessible chromatin elements. A) Workflow for human neocortical epigenetic characterization. See Materials and Methods for details. B-D) High-quality nuclei (2858 from 14 specimens) visualized by *t*-SNE and colored according to mapped transcriptomic cell types grouped into cell type subclass (B), sort strategy (C), or specimen (D). E) Transcriptomic abundances of eleven cell subclass-specific marker genes (median counts per million (CPM) within cell type) for 75 cell types identified in human MTG (Hodge et al., 2019). F) Eleven example subclass-specific marker genes demonstrating uniquely accessible chromatin elements in their vicinity (less than 50 kb distance to gene). Pileup heights are scaled proportionally to read number, and yellow bars highlight subclass-specific peaks for visualization. Dashed lines: introns, thick bars: exons, arrows: direction to gene body.

To identify putative regulatory elements within each subclass, we aggregated the data for all nuclei within each subclass, and identified peaks (median 411 bp across subclasses) using the Homer peak calling program (Heinz et al., 2010). This analysis revealed peaks proximal to recently identified transcriptomic subclass-specific marker genes (Hodge et al., 2019), further confirming our clustering and mapping strategy (Fig. 1E-F, S4, S5). We then used chromVAR (Schep et al., 2017) to identify differentially enriched transcription factor (TF) family motifs for known neuronal regulators. These TF motifs were strongly correlated with their TF transcript abundances from snRNA-seq data (Fig. S6A-H, Hodge et al., 2019). Together, these analyses demonstrated strong concordance between snRNA-seq and snATAC-seq data modalities at the cell subclass level.

### Concordance of epigenetic marks in human neurons from distinct profiling techniques

To assess the correspondence between ATAC-seq peaks and other methods of genomic element characterization, we first calculated the overlap between subclass snATAC-seq peaks and differentially methylated regions (DMRs) previously identified from human frontal cortex methylcytosine sequencing (Fig. S7, Lister et al., 2013; Luo et al., 2017). For every cell subclass, we observed a greater overlap of snATAC-seq peaks with DMRs than expected by chance (Fig. S6I), revealing thousands of neocortical regulatory elements (1,253 in microglia, to 123,665 in L23 neurons) by the intersection of both DMR and snATAC-seq data. In total, 27 ± 20% (mean ± sd) of all human peaks were also identified as DMRs. Peaks from all subclasses displayed greater than random conservation of primary DNA sequence as measured by phyloP scores (Fig. S6J, Pollard et al., 2010). Together, these analyses suggest that snATAC-seq faithfully detects DNA elements that have undergone positive selection through evolution, and likely play a functional role in these diverse cell types.

### Species-conserved and -divergent functional genomic elements

To identify regions of chromatin accessibility shared with mouse (“conserved”), as well as those present only in human or mouse (“divergent”), we aggregated mouse scATAC-seq peaks (Graybuck et al., 2019) to match our human dataset, and then computed Jaccard similarity coefficients between human and mouse subclasses by counting peak overlaps (Methods). All mouse subclasses displayed the highest similarity to the orthologous human subclasses, and all but one human subclass, hL56IT, matched reciprocally (Fig. 2A). Non-neuronal classes displayed the strongest cross-species similarity, followed by inhibitory neurons, whereas excitatory neurons displayed the weakest correspondence (Fig. 2A). The weak correspondence of excitatory neurons was likely partially due to regional mismatch between the mouse (V1) and human (MTG) sc/snATAC-seq datasets (Graybuck et al., 2019), since excitatory cortical neurons are known to be distinct across regions (Tasic et al., 2018). Nevertheless, this analysis yielded many more conserved peaks than expected by chance alone (Fig. 2B, ** FDR < 0.01 in each subclass). In sum, 34 ± 10% (mean ± sd) of all human peaks were also detected in matching mouse subclasses. Conserved peaks exhibited significantly greater primary sequence conservation than divergent peaks in both species (heteroscedastic t-test; human *t* = 10.3, p < 0.001; mouse *t* = 6.6, p < 0.001; Fig. 2C), supporting the notion that snATAC-seq reveals genomic elements that perform evolutionarily conserved functions. Consistent with this idea, using Linkage Disequilibrium Score Correlation (LDSC, Bulik-Sullivan et al., 2015; Finucane et al., 2015) we found that SNPs linked to educational attainment and schizophrenia were more closely associated with conserved neuronal peaks than with divergent neuronal peaks (Fig. S8A-C, see Materials and Methods for details). However, a notable counterpoint is the association between microglia and Alzheimer’s disease (Cusanovich et al., 2018; Girdhar et al., 2018; Skene et al., 2018; Nott et al., 2019) which showed stronger association in divergent human peaks than with conserved peaks (Fig. S8C), suggesting that Alzheimer’s-related microglial dysfunction is associated with human regulatory domains not present in mice (Zhou et al., 2020).

Additionally, we sought to understand how global genetic regulation differs across species and among cell subclasses. To address this, we first performed unbiased *de novo* identification of DNA sequence element motifs using MEME-CHIP (Bailey et al., 2009), which were then filtered for expression of a possible binding site-correlated transcription factor by RNA-seq (Fig. S9A, Tasic et al., 2018; Hodge et al., 2019). This analysis revealed several known cell subclass-specific TFs (for example SPI1/PU.1 in microglia and OLIG2 in oligodendrocytes/OPCs), and many unappreciated subclass-specific TFs (for example the TEAD motif is the most significant motif observed in human astrocytes but absent from mouse astrocytes [Fig. S9A]). Second, we wished to understand why peaks were present in common genomic repetitive elements. We noticed that, across cell subclasses and across species, divergent peaks more commonly overlapped with mobile repetitive genetic elements than conserved peaks do (Fig. S9B-C), suggesting a means for their dispersal, duplication, and mutagenesis during mammalian evolution (Hof et al., 2016; Gao et al., 2018). As a whole, these comparative analyses of single-cell epigenetic data furnish a wealth of knowledge about cell type identity determinants and origins.

**Figure 2.**
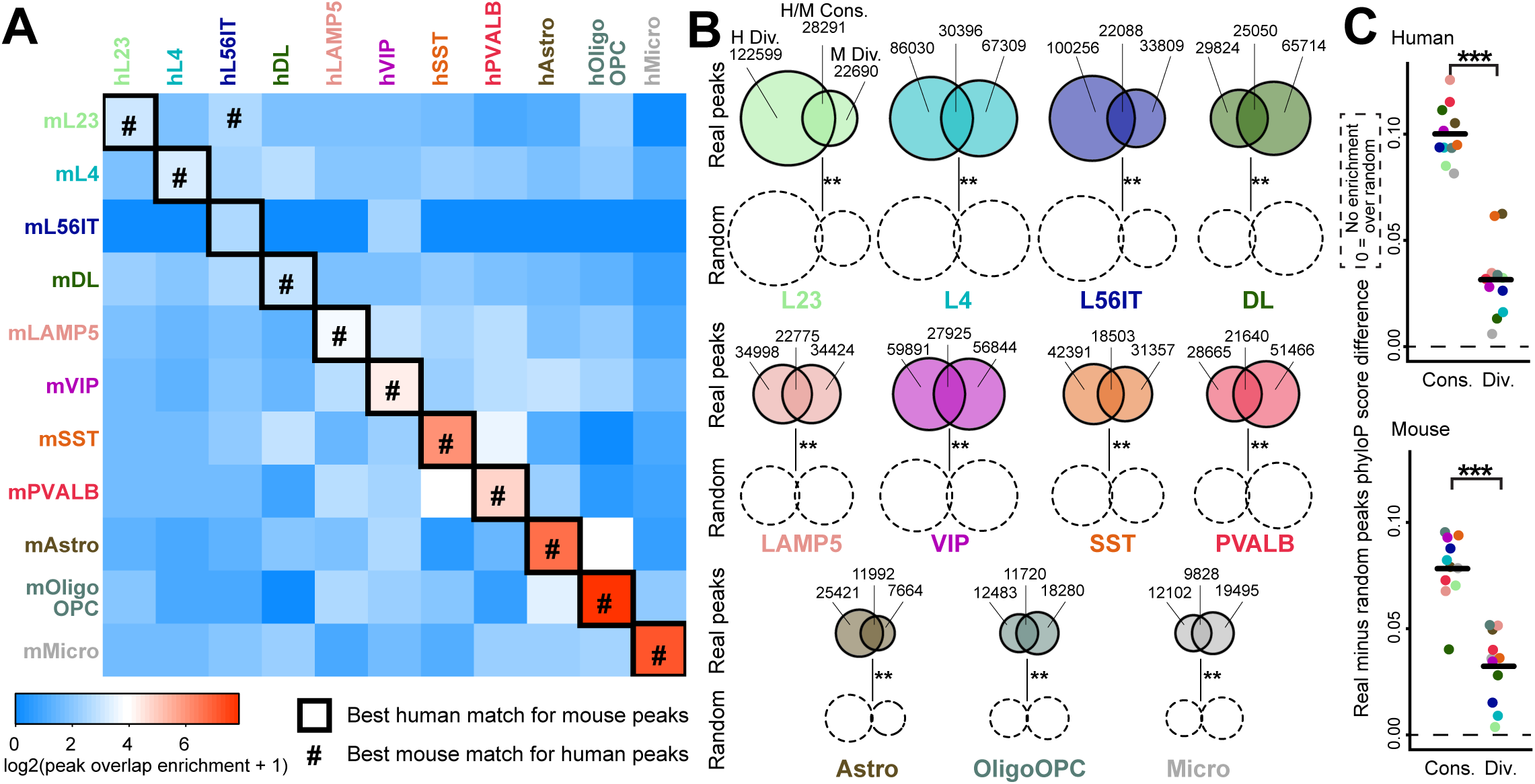
High conservation of human neocortical accessible genomic elements. A) Jaccard similarity coefficient enrichments (ratio of real to randomized peak positions) between human and mouse neocortical cell subclasses. Subclass-specific peaksets almost always best match their orthologous peakset across species. B) Visualization of conserved (Cons.) and divergent (Div.) peak counts across cell subclass in human and mouse. Conserved peaks are more frequent than expected by chance (** FDR < 0.01). C) Greater primary sequence conservation for conservedly accessible peaks than for divergently accessible peaks in both human and mouse. ***p < 0.001, by heteroscedastic t-test (human *t* = 10.3, df = 18.5; mouse *t* = 6.6, df = 19.9). Dashed line indicates no difference between real and randomized peak positions.

### Identifying functional enhancers using AAV reporter vectors

To determine whether ATAC-seq peaks might provide useful enhancers for developing novel genetic tools as had been previously shown (Dimidschstein et al., 2016; Nair et al., 2020) we cloned several peaks into a reporter AAV vector backbone and packaged viral particles with the mouse blood-brain barrier-penetrant capsid PHP.eB (Fig. 3A, Chan et al., 2017). We found that 1×1011 vg of AAV2/PHP.eB vector delivered intravenously (retro-orbital injection) demonstrated wide tropism for many brain neurons, as shown using the pan-neuronal promoter hSyn1 (Fig. 3B, McLean et al., 2014). Furthermore, we could also drive reporter expression in specific brain regions and neuron subclasses using enhancers; for example telencephalic interneurons with hDLXI56i (Fig. 3C, Zerucha et al., 2000; Dimidschstein et al., 2016).

We took several strategies to identify new enhancers with cell class and subclass-specific activity. In one approach, we manually identified peaks in the locus of known subclass-specific marker genes from snRNA-seq (Hodge et al., 2019), as shown for eHGT_078h, 058h, hDLXI56i (previously known), 019h, and 017h (Fig. 3D). We selected these peaks from the bulk population layer-specific open chromatin data before the single cell-resolution open chromatin data was available, based on enrichment in NeuN+ neurons over NeuN-glial cells and layer prevalence of the targeted cell subclass (Fig. S2; Hodge et al., 2019). In a second approach, we identified subclass-specific peaks that were conserved or divergent across human and mouse sn/scATAC-seq and DMR data (eg. eHGT_0128h, Fig. 3D, Luo et al., 2017). All enhancer-AAV vectors were then systemically delivered and validated for specific expression by both multiplexed FISH with hybridization chain reaction detection (mFISH-HCR, Choi et al., 2018), and by scRNA-seq from the visual cortex (Fig. 3E-G, Fig. S10) (Tasic et al., 2016, 2018).

We discovered several enhancer-AAV vectors that drove distinct reporter expression patterns consistent with their accessibility profiles (Fig. 3D-E). These expression patterns included pan-excitatory (eHGT_078h) and pan-inhibitory neocortical neurons (hDLXI56i, Zerucha et al., 2000; Dimidschstein et al., 2016), and subclass-specific *Rorb+* L4 and L56IT neocortical excitatory cells (eHGT_058h), and LAMP5 (eHGT_019h), SST and VIP (eHGT_017h), and PVALB (eHGT_128h) neocortical inhibitory cells. Multiplexed FISH-HCR demonstrated enrichment for the targeted cell subclasses with these enhancer-AAV vectors in V1, for example, 68 ± 9% of *Lamp5^+^* interneurons were labeled by eHGT_019h, and 82 ± 1% of *Slc17a7^+^Rorb^+^* L4 and L56IT neurons labeled by eHGT_058h (Fig. 3F). Finally, scRNA-seq confirmed the transcriptomic identity of the labeled cells with each of these viral vectors at the subclass (Fig. 3G) and type levels (Fig. S10).

### A collection of Parvalbumin-expressing (PVALB) neuron enhancer-AAVs

We sought to identify a collection of enhancers to enable access to PVALB interneurons which are important for cortical microcircuit regulation and can be dysfunctional in epilepsy, schizophrenia and Alzheimer’s disease (Cheah et al., 2012; Verret et al., 2012; Mukherjee et al., 2019). Since we identified many PVALB-specific open chromatin regions, we wanted to test how many could confer PVALB subclass-specific expression from AAV vectors. Importantly, there are currently few vectors specific for the PVALB subclass interneurons, especially those with validated expression in primates (Mehta et al., 2019; Vormstein-Schneider et al., 2020). We identified, cloned, and tested nineteen independent enhancers that show differing levels of specific accessibility for PVALB neocortical inhibitory cells (Fig. 4A). The first eight enhancers of the collection were selected using the strategy of identifying neuronal-enriched open chromatin regions near marker genes from layer-microdissected bulk population ATAC-seq data. Retrospective assessment of the snATAC-seq data showed none of these enhancers was specific for the PVALB subclass, and only two showed strong accessibility in PVALB cells (Fig. 4A). Only one enhancer, eHGT_023h, demonstrated reporter expression enriched in PVALB cells (Fig. 4A-B). Of the remaining eleven vectors that were selected based on single cell-resolution open chromatin enrichment in PVALB cells, five exhibited high selectivity for PVALB or PVALB/SST neocortical neurons in mouse as predicted from the human open chromatin data. Characterizing these enhancer-AAV vectors using both mFISH-HCR and scRNA-seq revealed some PVALB enhancers were less specific and some were highly specific (Fig. 4B-M). Neocortical V1 cells labeled by eHGT_023h were 47 ± 4% *Pvalb^+^* interneurons (Fig. 4B,H), whereas neocortical V1 cells labeled by eHGT_079h, eHGT_082h, eHGT_128h, and eHGT_140h were much more likely to be *Pvalb^+^* interneurons (95%-98% cells expressed *Pvalb* mRNA; Fig. 4D-G,J-M). Intermediate to these is eHGT_064h that labels both *Pvalb^+^* (50 ± 6%) and *Sst+* neurons in V1 (54 ± 1%, Fig. 4C,I), suggesting it enhances the nearby gene *CRHBP* which is specifically expressed in MGE-derived PVALB and SST interneurons (Tasic et al., 2018; Hodge et al., 2019). In agreement, 99% (183/185) of eHGT_064h-labeled cells expressed the MGE-derived inhibitory neuron marker *Lhx6*. Overall 6/19 (32%) of the tested enhancers showed PVALB-specific expression patterns, including 5/11 (45%) of those chosen with single cell-resolution epigenetics. These results not only reveal a new collection of human enhancer-based AAVs capable of marking PVALB or PVALB/SST subclass(es), they highlight the importance of high quality single-cell epigenetic data for efficient enhancer discovery.

Although some single cells marked by these PVALB-specific enhancers mapped to SST cell types, those SST cell types mostly expressed *Pvalb*, and as a result these vectors show greater specificity for *Pvalb*-expressing cells than for PVALB subclass cell types *per se* (Fig. 4I-M). We were surprised to find that the least specific PVALB enhancer eHGT_023h is located within an intron of *PVALB* itself, while the most specific PVALB enhancer eHGT_140h is not in the proximity of any known PVALB marker gene, and is instead in an intron of *NRF1*, a gene expressed non-specifically in most cell types of the neocortex. This highlights the importance of genome-wide enhancer discovery and demonstrates that restriction of the enhancer search to known marker genes may not support comprehensive development of the most specific viral tools. Most surprisingly, eHGT_079h, eHGT_128h and eHGT_140h were highly specific for mouse PVALB cells despite being accessible in human but not mouse PVALB cells, showing that the human enhancer sequence is sufficient to confer specificity even in a species that does not use that particular enhancer.

*PVALB/Pvalb-*expressing neurons are not just in neocortex but are also located throughout the CNS (Fig. 4N). We observed that some PVALB enhancer-AAV vectors labeled cells in known regions of *Pvalb* expression outside of neocortex (Fig. 4O-T). For example, eHGT_023h labeled Purkinje cells, mid/hindbrain nuclei, hippocampus and main olfactory bulb neurons (Fig. 4O,U), similar to *Pvalb* mRNA expression. Similarly, eHGT_082h labeled midbrain structures and deep cerebellar nuclei and main olfactory bulb, but not Purkinje cells (Fig. 4R,V). In contrast, eHGT_079h and eHGT_140h labeled mostly neocortical interneurons (Fig. 4Q,T). These results suggest our identified enhancer elements contain not only subtype targeting specificity, but also regional targeting selectivity which our selection strategy did not take into account.

### A collection of LAMP5 inhibitory subclass-specific enhancer-AAV vectors

In addition to PVALB-targeting enhancer-AAV vectors, we also discovered a group of four LAMP5 inhibitory cell-targeting vectors that show specificity for LAMP5 inhibitory neurons (eHGT_019h, 025h, 096h, and 098h, Fig. S11). Like for the PVALB collection, we identified these enhancers efficiently (4 of 12 enhancer-AAV vectors screened, 33% success rate, Fig. S11A), which we confirmed for cell subclass specificity by both mFISH-HCR (Fig. S11B-E) and by scRNA-seq (Fig. S10, S11F-I). Like the PVALB vectors, some of these enhancers (eHGT_025h and eHGT_096h) are cortex-specific, and in contrast some (eHGT_019h and eHGT_098h) show subcortical expression domains which are shared with the *Lamp5* expression pattern (Fig. S11J-N). All vectors were highly enriched for rare *Gad1*-expressing inhibitory cells, despite most *Lamp5*+ cells in the cortex belonging to the excitatory class. Together the PVALB and LAMP5 enhancers suggest that snATAC-seq data can efficiently identify enhancer collections for multiple target neuronal populations.

### Enhancer AAV vectors enable genetic access to NHP and human neocortical cell types

Our goal is to develop AAV tools that retain subclass selectivity across species and especially in human. To first test if cell class-selective expression was maintained in NHP neocortex, we injected our enhancer-AAV vectors intraparenchymally in multiple sites in one *M. mulatta* occipital cortex and then evaluated expression selectivity after 50 days (Fig. 5A). We tested three PVALB-specific vectors identified from our mouse primary screen in NHP (eHGT_079h, 128h, and 140h) (Fig. 5B-D). Immunohistochemistry showed that vast majority of vector-labeled cells expressed PVALB (86, 95, and 95% of SYFP2+ cells, respectively). Furthermore, nearly all the PVALB^+^ cells throughout the cortical column expressed SYFP2 from the vectors eHGT_128h and 140h (89 and 92%), whereas eHGT_079h showed fewer labeled PVALB^+^ cells (62%). This finding indicates not only that these vectors are highly specific for primate neocortical PVALB cells, but also PVALB cell labeling can be nearly complete. We also injected eHGT_140h into this animal’s temporal cortex which again showed high labeling specificity (82%) and completeness (76%), demonstrating that this enhancer-AAV vector functions across cortical regions in NHP.

To improve the throughput of NHP testing and to extend testing to human tissue, we leveraged a slice culture platform for testing enhancer-AAVs. We obtained NHP MTG tissue from regularly scheduled *M. nemestrina* harvests at the Washington National Primate Center (Fig. 5E). Virally transduced *ex vivo* slices cultured for one to two showed selective cell distributions and morphologies consistent with the mouse observations, including eHGT_078m (the mouse ortholog of eHGT_078h) in L2-6 excitatory neurons (Fig. 5F), and eHGT_058h in L3-5 pyramidal neurons (Fig. 5G). We boosted expression of hDLXI56i by generating 3x(hDLXI56i core), containing three core elements of hDLXI56i in tandem array, which resulted in bright expression in non-pyramidal neurons (Fig. 5H). We confirmed high specificity of labeling in these cultures by mFISH-HCR (Fig. 5I-K), which demonstrated a similar high specificity of these three enhancers in NHP *ex vivo* slices as that seen in mouse (compare Figs. 3E-G and 5I-K). However, certain vectors including the PVALB vectors eHGT_079h, 128h, and 140h displayed substantially reduced specificity in NHP slice cultures relative to that seen in mouse and NHP *in vivo* (compare Figs. 4D,F,G,J,L,M and 5B-D and 5L-N). This loss of specificity was particularly profound in the case of eHGT_140h (mouse *in vivo* 98-99%, NHP *in vivo* 82-95%, and NHP *ex vivo* 7% specific, Figs. 4G,M and 5D and 5N). These data suggest that some cell subclass-specific enhancers change activity in slice culture context, whereas others maintain their specific mode of action. Further studies will be required to establish why certain vectors lose specificity of expression in NHP slice culture.

To establish the specificity of enhancer-AAV vectors in human tissue directly, we first tested whether hDLXI56i-AAV vectors would label forebrain inhibitory neurons in human *ex vivo* brain slice cultures (Fig. 6A, Ting et al., 2018), following prior demonstration of inhibitory cell specificity in human stem cell-derived neurons *in vitro* (Dimidschstein et al., 2016). We observed that hDLXI56i labeled neurons of non-pyramidal morphologies throughout the neocortical layers (Fig. 6B), and profiling these cells by scRNA-seq revealed that the great majority of them are *GAD1+* inhibitory neurons (97%) of multiple transcriptomic cell types (Fig. 6C). Next we sought to find new enhancers for increased control of targeting human neocortical subclasses. We selected a single region containing a conserved regulatory element found near the CGE-derived LAMP5/VIP subclass marker gene *CXCL14* (region eHGT_022, Fig. 6D). AAV vectors containing either the human or mouse ortholog of eHGT_022 (eHGT_022h and eHGT_022m, respectively) were able to drive expression in upper-layer-enriched interneurons in both mouse and human (via live human *ex vivo* slice culture infection, Fig. 6E). When profiled by immunohistochemistry or scRNA-seq, reporter-positive cells in both mouse and human mapped specifically to LAMP5 and VIP inhibitory neurons (Fig. 6F-G). eHGT_022h-labeled mouse cells were 95 ± 7% Lamp5^+^ or VIP^+^ by IHC or 99% (122/123) Lamp5^+^ or VIP^+^ by scRNA-seq; eHGT_022m-labeled mouse cells were 95 ± 3% Lamp5^+^ or VIP^+^ by IHC; and eHGT_022h-labeled human cells were 95% (101/106) Lamp5^+^ or VIP^+^ by scRNA-seq. Thus, the eHGT_022h/m enhancer-AAV vectors mark CGE-derived inhibitory cells across species.

**Figure 3.**
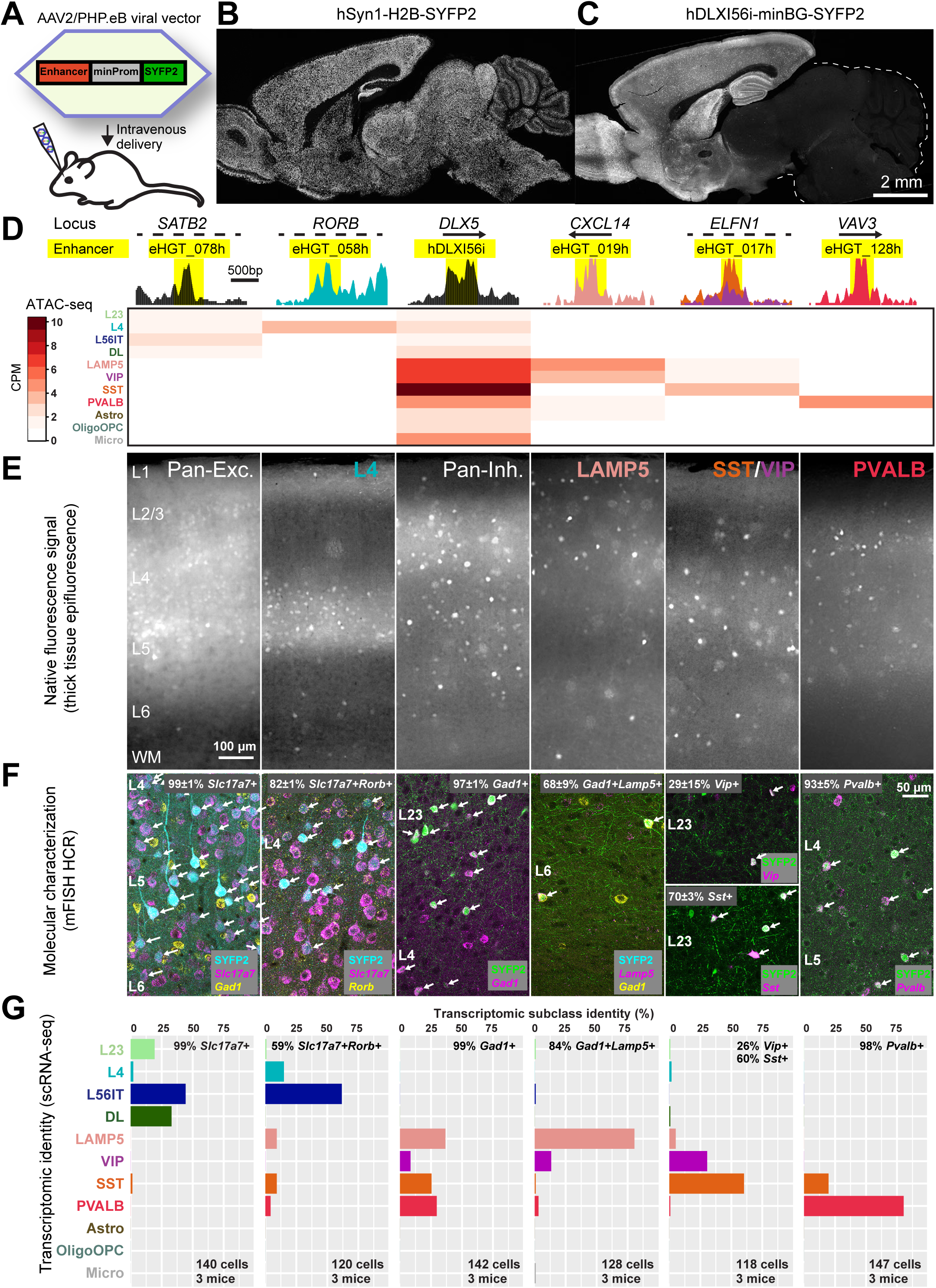
Accessible chromatin elements furnish cell subclass-specific AAV genetic tools. A) AAV2/PHP.eB viral reporter vector design for testing presumptive enhancers cloned upstream of a minimal promoter and SYFP2 reporter expression cassette in mouse retro-orbital assay. B) Transgene expression from AAV-hSyn1-H2B-SYFP2 in most neurons throughout mouse brain. C) Transgene expression from AAV-hDLXI56i-minBG-SYFP2 in mouse forebrain interneurons, in agreement with previous reports (Zerucha et al., 2000; Dimidschstein et al., 2016). D) Several identified enhancers showing ATAC-seq peaks in distinct target cell subclasses. Each selected enhancer is highlighted in yellow on read pileups, and heatmap below demonstrates ATAC-seq read CPM in all cell subclasses. E) Distinct expression patterns from these enhancer-AAV vectors in live 300 micron-thick slices of primary visual cortex (V1) after retro-orbital delivery, consistent with different subclass-specific expression patterns. F) Multiplexed FISH-HCR in V1 region revealing differing subclass specificities from various enhancer-AAV vectors. Text represents mean ± standard deviation for labeling specificity across 3 independent mice. G) Single cell RNA-seq on sorted individual SYFP2^+^ cells from V1 region confirming distinct cell subclass transcriptomic identities labeled by the highlighted enhancer-AAV vectors.

**Figure 4.**
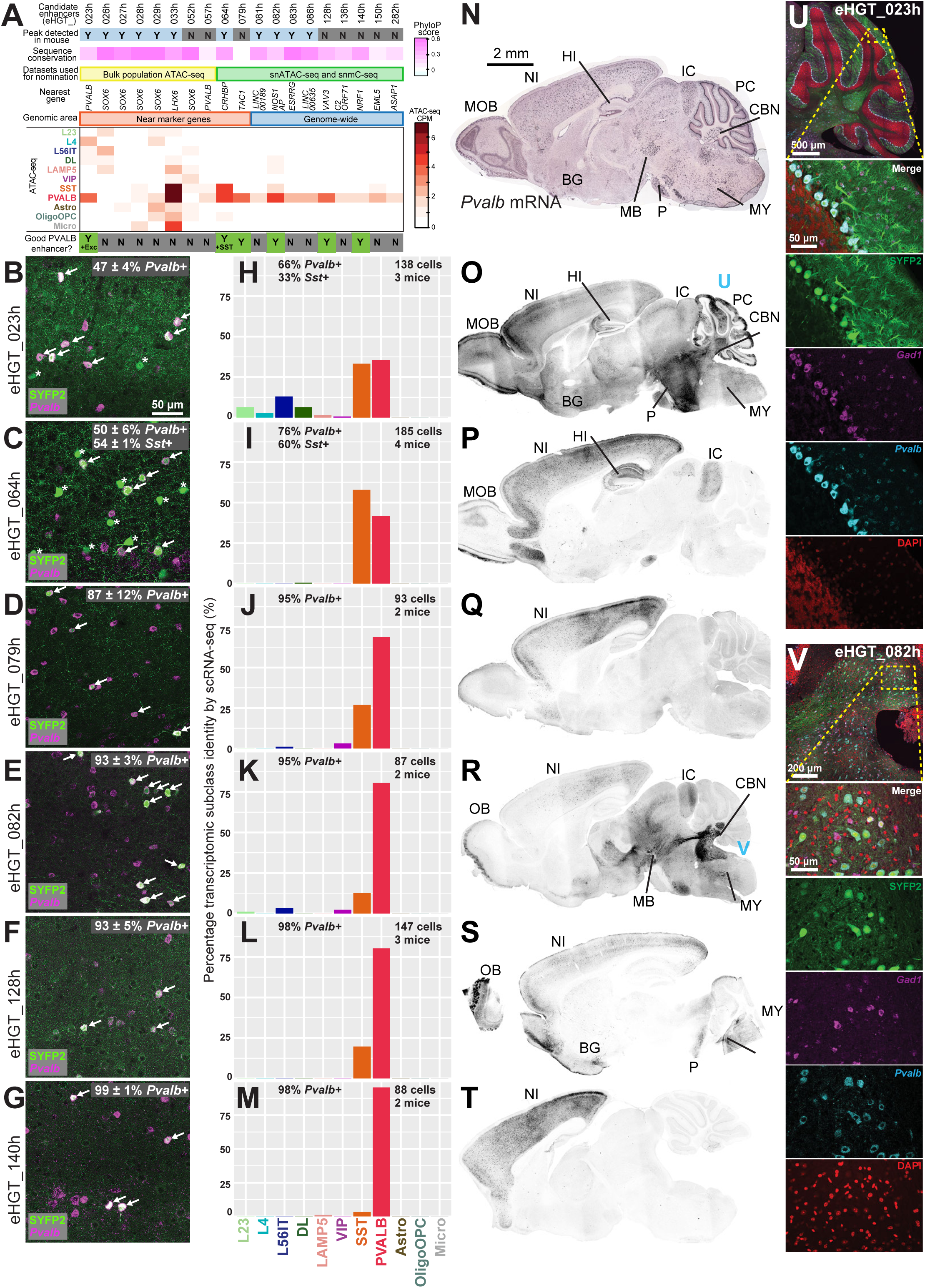
PVALB neocortical interneuron enhancers display distinct subcortical expression patterns. A) Nineteen putative PVALB enhancers from snATAC-seq data cloned into AAV vectors. Six of the nineteen (32%) exhibited high selectivity for PVALB cells in mouse retro-orbital assay (indicated with green boxes). B-G) mFISH HCR in L2/3 of V1 demonstrating positive labeling of *Pvalb^+^* cells (arrows) by each of the indicated enhancer-AAV vectors. eHGT_023h and eHGT_064h also label non-*Pvalb^+^* cells (asterisks). Percentages indicate the mean specificity ± standard deviation of SYFP2 labeling for *Pvalb*^+^ cells across 3 independent mice. H-M) Single-cell RNA-seq in V1 confirming the PVALB transcriptomic cell subclass identity of enhancer-AAV vector-labeled cells. Bargraph shows the percentage of single cells that map to a transcriptomic cell type within that subclass. In contrast, the percentages given in the text are the percentage of cells recovered that expressed the indicated gene. Note that although only 65% of the eHGT_079h-marked cell types mapped to the PVALB subclass, 94% of the eHGT_079h-marked cells expressed *Pvalb* mRNA. This is because several SST subclass cell types also express *Pvalb* mRNA. N) *Pvalb* mRNA expression pattern (Allen Institute public ISH data) with multiple sites of expression throughout mouse brain. O-T) Labeling of both neocortical PVALB cells, and also various subcortical brain regions, by PVALB-selective enhancers. These subcortical brain regions are also seen in the endogenous *Pvalb* mRNA expression pattern. Two enhancers (eHGT_079h and 140h) show exceptional specificity to neocortical PVALB cells. Abbreviations: NI neocortical inhibitory neurons, HI hippocampal inhibitory neurons, MOB main olfactory bulb, BG basal ganglia, MB midbrain nuclei, MY medulla nuclei, P pons, IC inferior colliculus, PC Purkinje cells, CBN deep cerebellar nuclei. U-V) Subcortical labeling by eHGT_023h in Purkinje cells (U) and by eHGT_082h in CBN (V). eHGT_023h-labeled Purkinje cells are *Pvalb^+^Gad1+*, and eHGT_082h-labeled CBN cells are either *Pvalb^+^Gad1+* or *Pvalb^+^Gad1-*.

**Figure 5.**
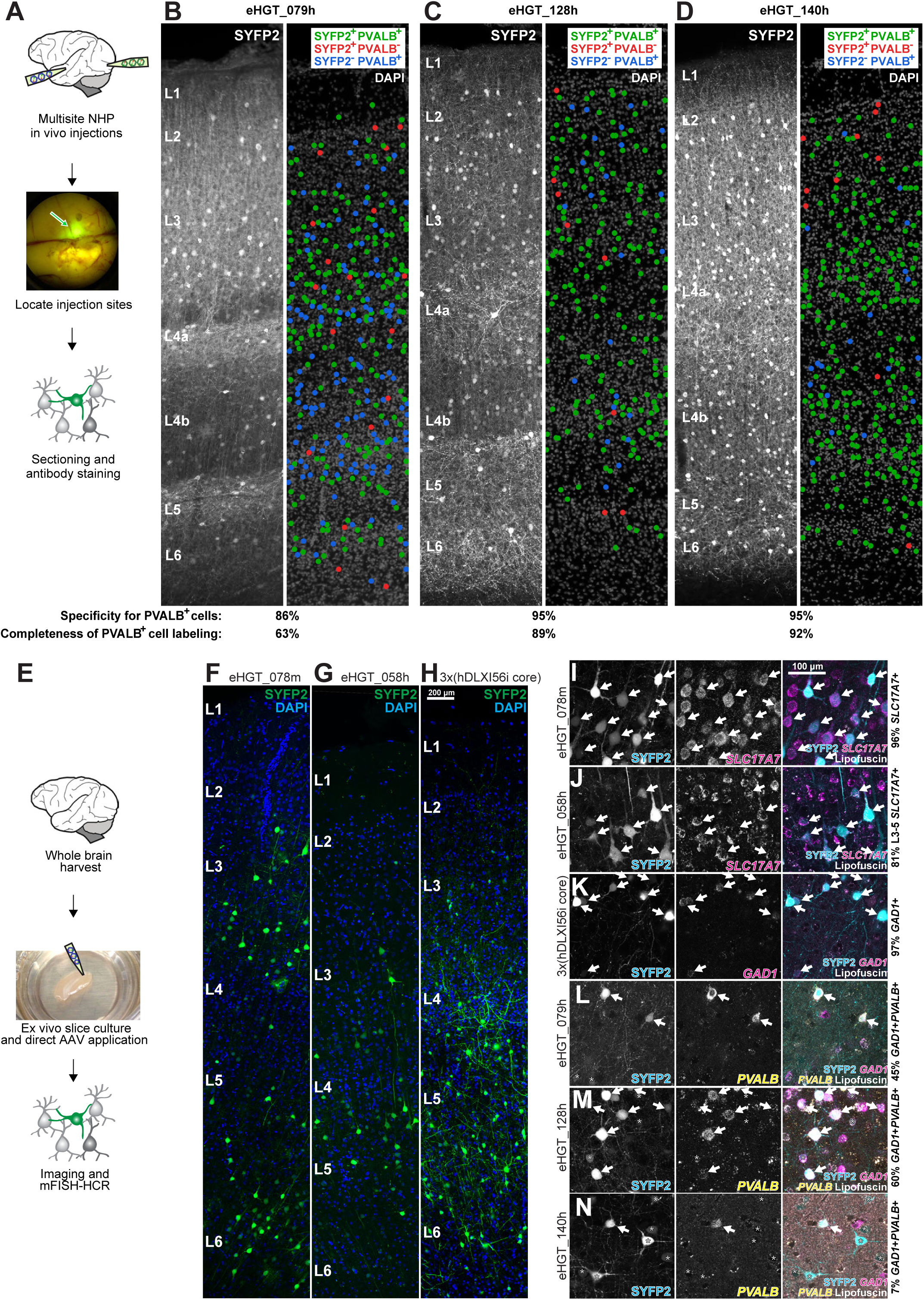
Multiple enhancer vectors demonstrate cell subclass specificity in NHP neocortical tissue. A) Workflow for *in vivo* AAV vector testing by intraparenchymal injection in NHP brain. B-D) Injection of eHGT_079h, 128h, and 140h AAV vectors into NHP occipital cortex. These three vectors label PVALB cells throughout the cortical column with high specificity. eHGT_128h and 140h label nearly all PVALB cells, but eHGT_079h labels fewer PVALB cells. Dots indicate the positions of counted cells observed by coimmunostaining with anti-GFP and anti-PVALB antibodies (>300 cells were counted per vector in one experiment). E) Workflow for acquiring fresh NHP neocortical tissue for AAV vector testing *ex vivo*. F-H) Transduction of *ex vivo* NHP neocortical tissue with various AAV2/PHP.eB enhancer-reporter vectors, resulting in diverse expression patterns. eHGT_078m labels excitatory neurons throughout all layers (F), eHGT_058h labels excitatory neurons primarily in L3-4 (G), and 3x(hDLXI56i core) labels inhibitory neurons (H). I-N) NHP neocortical cell subclass specificity of AAV vector labeling confirmed by mFISH-HCR. eHGT_078m, 058h, and 3x(hDLXI56i core) demonstrate high specificity similar to that seen in mouse retro-orbital assay, but eHGT_079h, 128h, and 140h show reduced specificity compared to that seen in mouse retro-orbital assay and NHP *in vivo* assay. Arrows highlight specifically labeled on-target cell types, and asterisks mark off-target labeled cells. Text represents mean for labeling specificity across 1-2 independent transduction experiments (>100 cells counted per vector per experiment).

**Figure 6.**
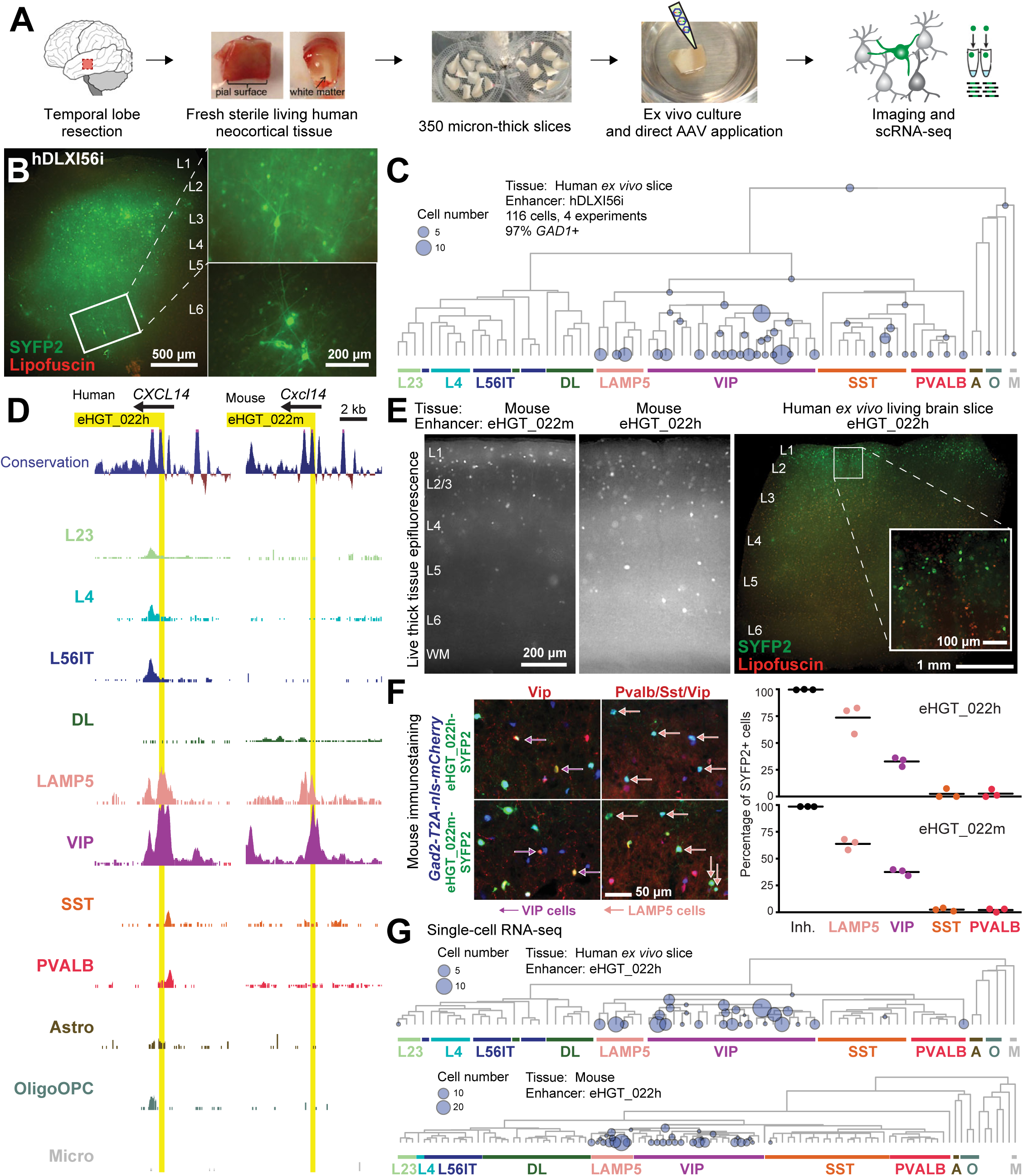
eHGT_022h/m enables non-species-restricted tools for genetic access of CGE-derived LAMP5 and VIP neurons in human brain tissue. A) Workflow for acquiring fresh human neurosurgical tissue for AAV vector testing. B) AAV-hDLXI56i-minBG-SYFP2 transduction of human *ex vivo* brain slices. This vector labels scattered neurons with diverse non-pyramidal cellular morphologies spanning all neocortical layers (live thick tissue epifluorescence imaging). C) Molecular identity of AAV-hDLXI56i-minBG-SYFP2+ human cells by scRNA-seq. The great majority of AAV-hDLXI56i-minBG-SYFP2+ human cells are inhibitory neurons of multiple transcriptomic types. Dendrogram represents human MTG taxonomy (Hodge et al., 2019), leaves represent 75 transcriptomic cell types, and circles represent numbers of cell types detected. Circles on intermediate nodes of the dendrogram represent incomplete mapping to cell type. Data from 4 independent experiments are shown. D) eHGT_022h/m accessibility only in LAMP5 and VIP neuron subclasses in both human and mouse. E) eHGT_022h/m expression in primarily upper-layer scattered neurons in mouse V1 and in human *ex vivo* neocortical brain slice cultures. F) Immunostaining eHGT_022h/m-SYFP2+ V1 mouse cells suggesting they are primarily *VIP^+^* or *Lamp5*+ (Gad2+Pvalb-Sst-Vip-) neurons. Over 1000 cells from 3 mice counted for each reporter. G) Molecular identity of eHGT_022h-SYFP2+ cells by scRNA-seq. The great majority of eHGT_022h-SYFP2+ cells from mouse (123 cells from three independent mice) and human (106 cells from one experiment) are VIP and LAMP5 neurons. Dendrogram leaves represent transcriptomic cell types (75 in human and 111 in mouse V1, Tasic et al., 2018; Hodge et al., 2019) and circle sizes represent number of cells detected at each node.

## Discussion

Here we present data and methodology to generate and evaluate AAV-based viral tools that drive brain cell subclass-specific transgene expression from mouse to primate. First, we report and characterize a subclass-resolution snATAC-seq dataset of human neocortex. Second, we compare this human open chromatin dataset to a comparable mouse cortex dataset (Graybuck et al., 2019) to identify conserved and divergent subclass-selective putative enhancers. Third, we show that many enhancers yield subclass-selective expression in mice once inserted into an AAV vector upstream of a minimal promoter and reporter. Fourth, we present a collection of AAV vectors designed to target PVALB and LAMP5 inhibitory cells, with an efficient on-target rate of over 30% of tested vectors yielding the desired PVALB or LAMP5 selective expression in the cortex. Fifth, we demonstrate that many enhancers parcellate the expression patterns of targeted marker genes and show substantial regional variation, with some labeling multiple subcortical neuron populations and others highly specific for neocortical interneuron populations. And sixth, we confirm several of these AAV-reporters drive expression in primate *in vivo* and/or *ex vivo* cultured neocortical slices. In particular, eHGT_022h/m vectors label CGE-derived cortical inhibitory neuron subclasses in mouse and human, while eHGT_079h, 128h, and 140h label PVALB neocortical inhibitory cells in mouse and NHP. These AAV vectors constitute some of the first genetic tools with validated subclass selectivity for neocortical cell types across multiple mammalian species (see Fig. 7 for summary). We propose that snATAC-seq-assisted enhancer discovery is a generalizable strategy to efficiently identify cell subclass-specific enhancers (Graybuck et al., 2019) for the observation and perturbation of brain cell subclasses and types in a non-species-restricted manner.

**Figure 7.**
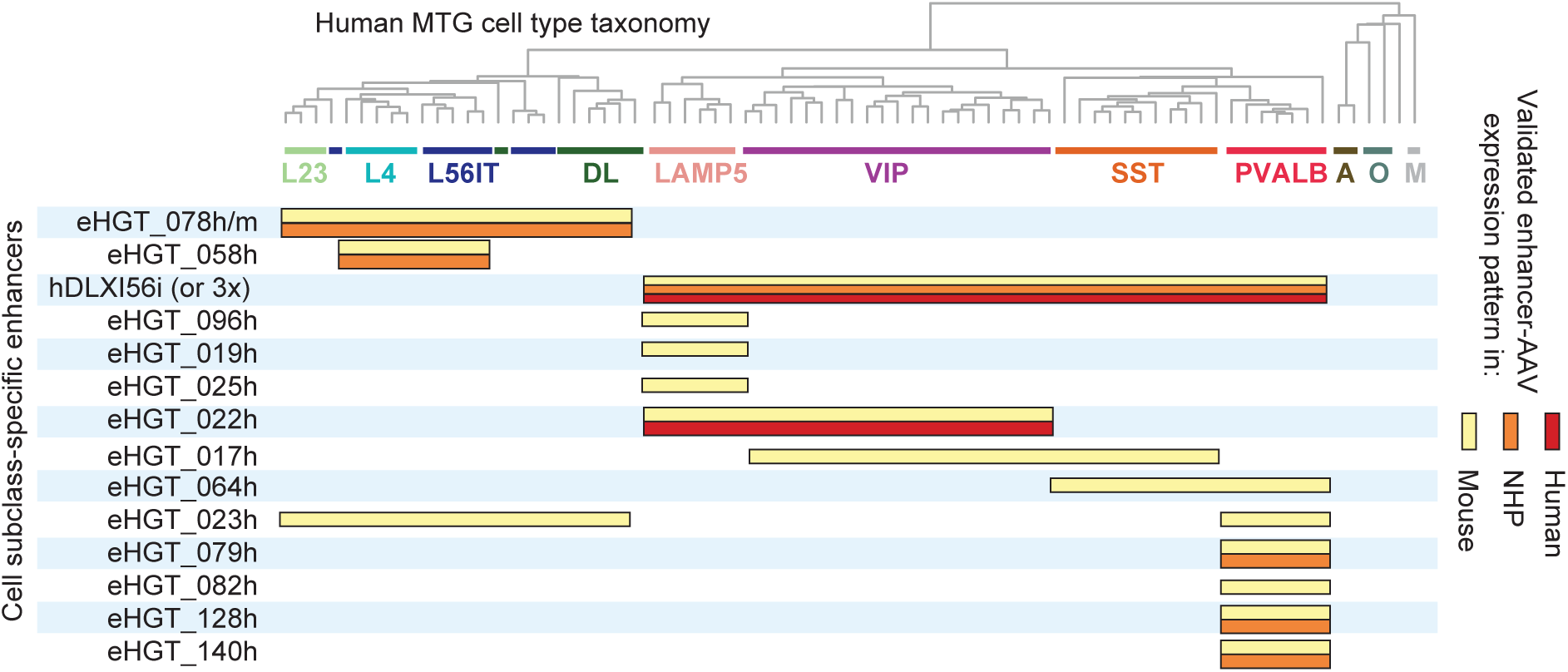
A growing catalog of well-validated, non-species-restricted AAV-based genetic tools for targeting neocortical cell subclasses and types. The Allen Institute Human Genetic Tools project is generating a catalog of validated enhancer-AAV vectors that permit targeting of major cell classes and subclasses in the neocortex. Bars indicate the cell classes and subclasses targeted by the indicated enhancer, as validated by mFISH-HCR and scRNA-seq in mouse and NHP and human neocortical tissue.

### High-quality and high-resolution epigenetic datasets uncover enhancers

Our subclass-resolution snATAC-seq profiling experiments from the human MTG were critical for the development of the new cell subclass-specific viral tools presented here. Most enhancers identified in our study were not visible in bulk epigenetic datasets (The ENCODE Project Consortium, 2012; Fullard et al., 2018). This is partially due to bulk data masking cell subclass specific peaks when the target cell population is not abundant. For instance, all inhibitory cells comprise only 20-30% of all cortical neurons and each inhibitory subclass is just a fraction of the total inhibitory cells (Tasic et al., 2018; Hodge et al., 2019). Also, many previous human datasets were prepared from frozen postmortem tissues, which yielded lower signal to noise data in our hands (data not shown). Single nuclear resolution ATAC-seq using freshly isolated nuclei from acutely resected neurosurgical tissue was key generate high quality data that enabled us to efficiently identify subclass-specific enhancers.

With high-quality and single cell-resolution epigenetic data across species we have discovered new insights into the gene expression regulatory apparatus, and how it varies across subclasses and species. First, ATAC-seq and single cell methylC studies frequently agree at the single cell level (27% of peaks detected as DMRs, Fig. S6, Luo et al., 2017) showing overlap of hypomethylated regions with open chromatin. Second, human and mouse orthologous neocortical cell peaks overlap strongly between species with 34% of human ATAC-seq peaks also detected in mouse (Fig. 2A-B). Third, we can predict transcription factors that are enriched at cell subclass- and species-specific peaks (Fig. S9A), observe the mechanisms by which enhancers might evolve and expand cell type diversity through colocalization with motile genetic elements (Fig. S9B-C), and ascertain how different cell subclass enhancers associate with neurological diseases (Fig. S8). In summary, high-quality epigenetic datasets enable biological discovery about the evolution, transcriptomic control and disease association of human brain cell subclasses.

### Building the next generation of AAV-based genetic tools

The identification of functional enhancers has been a challenge, and multiple prediction criteria have been used. Selection of open chromatin regions proximal to known marker genes is sometimes successful, but we favor a genome-wide and gene-agnostic selection criteria. The best proximal enhancer to the *PVALB* gene, eHGT_023h, was only weakly specific, while eHGT_082h, eHGT_128h and eHGT_140h are highly specific for PVALB cells but do not reside in the proximity of any known PVALB cell marker gene. Yet eHGT_082h drove specific reporter expression in both neocortical PVALB inhibitory cells and *Pvalb^+^* cells in the deep cerebellar nucleus. Importantly, removing the restriction of sampling open chromatin regions proximal to known marker genes greatly increases the number of specific elements available to test. We also noted that conservation of sequence or open chromatin across mouse and human isn’t essential to find functional enhancers that drive subclass selective expression in mouse and primate. For example, eHGT_079h showed weak sequence conservation and was only observed in the human snATAC-seq data, yet showed highly selective expression in PVALB subclass cells. This demonstrates that identification of conserved enhancers may not be necessary to identify cross-species-specific vectors, and that a human enhancer sequence can maintain specificity even in a species that doesn’t use that particular enhancer. Future studies could reveal insights into the endogenous roles of these distal or non-conserved selective enhancers.

We have learned several lessons about enhancer discovery through our analysis of the AAV vectors presented in this study. Our technique for screening AAV vectors *in vivo* enabled us to see whole-brain expression patterns, and we were surprised by the diversity of subcortical cell populations that were labeled by our PVALB viral vectors. These vectors labeled the PVALB cell subclass within the neocortex as predicted from the epigenetic data, but their subcortical expression patterns vary dramatically. For some vectors the expression is neocortex-restricted, while other vectors also label various *Pvalb^+^* cell populations in sub-cortical regions. Since all epigenetic profiling that informed our enhancer discovery is from neocortex (mouse V1 [Graybuck et al., 2019], human MTG, or mouse or human frontal cortex [Luo et al., 2017]), we could not have predicted whether these enhancers would drive expression in other brain regions. We now anticipate single cell epigenetic profiling studies of multiple brain regions will be required to predict of regional selectivity of enhancer activity. Such analyses will be critical for gene therapy vectors that will be delivered systemically if transgene expression is to be restricted to the targeted region. Additionally, the distinct patterns of expression for our PVALB vectors hint towards an additive “Lego logic” of enhancers that must act together to yield a gene’s complete expression pattern. Individual enhancers, thus, often show a more restricted expression pattern than seen from the best marker genes, and could be a method to produce targeted expression of transgenes in discrete brain regions.

Massively parallel enhancer screens (Shen et al., 2016; Hrvatin et al., 2019) have identified few selective enhancers from libraries of hundreds or thousands of candidate enhancers. In contrast, we show the efficient identification of selective enhancers using a one-by-one enhancer screening strategy informed by single cell-resolution epigenetic data. Six of the 19 viral vectors tested for specificity in PVALB cells showed significant on-target reporter expression (and 4 of 12 for LAMP5 cells), with improved efficiency for PVALB-specific labeling for the subset of putative enhancers selected from the single cell epigenetic data (45% showed on-target expression). Since we identified tens of thousands of putative subclass-specific enhancers from our limited epigenetic data, we expect that we will be able to identify many additional viral vectors for each subclass without implementing more complex batched screening strategies. The primary limitation for our enhancer discovery is the resolution of available epigenetic datasets: as that improves, so too will our ability to predict highly specific putative enhancers—possibly at the granular level of cell types. Once enhancers are identified, a second limitation is rigorous validation experiments to characterize the expression patterns of these enhancer vectors, and optimization of their activity to improve their utility as research reagents for characterization of cell type function across species. Although we have been efficient in targeting PVALB and LAMP5 cells, it is possible that certain rare cell types may be difficult to target with enhancers, which could make batched screening essential, and/or require the implementation of intersectional labeling strategies.

*Enhancer-AAVs function across mammalian species.* Testing the function of enhancers in primates is essential to determine if the enhancer-AAVs will be functional across species (Jüttner et al., 2019; Mehta et al., 2019). Two primate preparations were used in this study: *in vivo* injections into NHP neocortex, and *ex vivo* organotypic cultured slices from NHP or human neocortex. We tested three top PVALB vectors using intraparenchymal injections into the occipital cortex of a rhesus macaque monkey and saw excellent conservation cell subclass-selective expression from mouse to monkey. This confirmed that our selection method and mouse screening strategy is effective for identifying enhancers that function across species.

Since such injections in NHP are costly, challenging to execute, cannot be effectively scaled, and cannot be applied in human, we also applied the approach of AAV transduction in *ex vivo* organotypic slice culture of primate brain tissue (Ting et al., 2018). Using this strategy, we were able to show that the expression from eHGT_022h was conserved in mouse and human, and that several other vectors showed similar subclass specificities between mouse *in vivo* and NHP *ex vivo*. However, some vectors such as eHGT_140h exhibited altered specificity of labeling in the ex vivo paradigm, demonstrating that some but not all enhancers can be screened using ex vivo slice cultures, and underscoring the importance of ultimate validation in vivo. To better understand the changes that occur in the *ex vivo* cultured cells, and to generate AAVs that also maintain specificity in culture, more extensive transcriptomic and epigenetic profiling of *ex vivo* cultured cells will be required.

### Conclusion

Human brain functions and diseases are often difficult to study because model organisms do not recapitulate human brain circuitry or display clear clinically relevant phenotypes. It is thus critical to understand human brain-specific circuit components and their regulatory apparatus, because this may provide avenues for therapeutic intervention. We have catalogued human neocortical chromatin accessibility with single cell resolution, which deepens our knowledge of human brain cell subclass-specific gene regulation. Guided by the epigenetic data, we have built a new collection of cell subclass-specific AAV tools to target these cell types, and established an efficient platform to validate enhancer-AAV activity across species. These AAV tools, screening platform and epigenetic knowledge will guide a new experimental understanding of brain circuitry function, highlight a process for developing new AAVs that drive selective gene expression in cell populations across the brain, and may enable more precise AAV-based gene therapy vectors for unmet clinical needs.

## Acknowledgements

We thank Allison Beller, Nathan Hansen, Caryl Tongco, Jae-Guen Yoon, and Gina DeNoble for assistance with obtaining patient consent and human neurosurgical tissue research specimens. We thank Rebecca D. Hodge and Trygve E. Bakken and Zizhen Yao for assistance with sc/snRNA-seq data. We thank Lisa McConnell for assisting with NHP virus injection surgery and also NHP animal care. We thank Allen Institute Tissue Procurement and Facilities teams for institutional support during tissue collections.

## Author contributions

JTT, NS, EEH, TC, ND, and JKM performed tissue processing and flow cytometry. EEH, BPL, and JKM performed ATAC-seq with assistance from DB and KAS. JKM analyzed ATAC-seq data using techniques developed by LTG, with assistance from SS, JAM, LTG, JG, and YD. BPL, JTT, and JKM curated candidate enhancers for testing. EEH, JKM, JTT, RAM, and XOA performed AAV vector design and molecular biology. JKM and JTT tested AAV vectors with assistance from PC and XOA. RPG, CC, JGO, ALK, CDK, and DLS procured human surgical tissue for research. JTT and JKM performed human *ex vivo* brain slice culture and viral labeling experiments with assistance from PC. JTM, LL and YD performed mFISH-HCR. Single cell RNA-seq was conducted by JKM, DB, KAS, and analysis by OF, JG, and BPL.

MM, SY, AC, EEH, and XOA performed viral packaging. JTT carried out non-human primate *ex vivo* slice culture experiments with assistance from GDH, NW, and XOA. GDH and YK performed NHP *in vivo* virus injection surgery with assistance from JTT. JKM and JTT processed NHP brain tissue from *in vivo* virus testing with assistance from VO and YB. BPL, JTT, JKM, and EL conceived of the study design. JKM wrote the manuscript and prepared figures. LTG and BT provided mouse ATAC-seq data. VG provided PHP.eB capsid plasmid DNA. SMS provided program and budgetary management. HZ and ESL provided program leadership.

## Data and materials availability

scATAC-seq and scRNA-seq data will be deposited to GEO or dbGAP. Software code used for data analysis and visualization is available from GitHub at https://github.com/AllenInstitute/graybuck2019analysis/.

## Supplementary Materials

Materials and Methods Figures S1-S11

Supplementary References

