## Supplementary Material for "Functional enhancer elements drive subclass-selective expression from mouse to primate neocortex"

Contents:

Materials and Methods

Supplementary References

Figures S1-S11

#### Materials and Methods

*Neurosurgical tissue acquisition.* We receive regular acute neurosurgical brain tissue donations at the Allen Institute for Brain Science. These samples are excised as a matter of course to access the epileptic focus or tumor. All samples used in this study were derived from temporal cortex, most frequently middle temporal gyrus (MTG). These samples are immersed in pre-carbogenated ACSF.7 (recipe below), transported to the Allen Institute for Brain Science rapidly with carbogenation, and sliced on a compresstome (Precisionary Instruments, Greenville NC USA, catalog #VF-200) into 350 µm slices, and continuously carbogented in ACSF.7 until dissociation.

*Bulk tissue ATAC-seq.* We harvested MTG tissue slices after carbogen bubbling in ACSF.7 for up to 16 hours, and we treated with NeuroTrace 500/525 (catalog # N21480 from ThermoFisher Scientific, 1/100 in ACSF.7) to highlight layered cortex structure. With fine forceps we trimmed away white matter and meningeal tissues, and then dissected layers 1-6 into six different low-binding Eppendorf 1.5 mL tubes (MilliporeSigma catalog # Z666548) under a fluorescence microscope as in Hodge et al. (Hodge et al., 2019). We discarded supernatant and replaced with 50-100 µL of Nextera DNA library reaction (#FC-121-1031 from Illumina) containing 0.1% IGEPAL-630 (NP-40 alternative), and then pipetted up and down vigorously 25-

50 times using a P200 pipette, and then incubated at 37˚C for one hour for transposition. We then added 1 mL of Homogenization Buffer (recipe below) to quench the reaction, pelleted samples at 1000g for 5 minutes at 4˚C, resuspended samples in 1 mL fresh homogenization buffer, released nuclei from samples using ~10-15 strokes of a loose-fitting dounce pestle followed by ~10-15 strokes of a tight-fitting dounce pestle, then filtered nuclei with a 70 µm nylon mesh strainer, and pelleted nuclei at 1,000xg for 10 minutes at 4˚C. To stain, we resuspended nuclei in 500 µL of ice-cold Blocking Buffer (recipe below) containing 1/500 PE-NeuN antibody (MilliporeSigma catalog # FCMAB317PE) and 1 µg/mL 4'-diamino-phenylindazole (DAPI, MilliporeSigma catalog # D9542), rocked samples for 30 minutes at 4˚C, then pelleted at 1,000xg for 5 minutes at 4˚C, and finally resuspended samples in 500 µL fresh ice-cold blocking buffer before sorting cells on a FacsARIA III.

Using scatter profiles to eliminate debris and doublets, we sorted bulk samples as $DAPI^+NeuN^+$ from layers 1-6, or as $DAPI^+NeuN^-$ from layer 1 and layer 5 samples, at 5,000-10,000 cells per sample, into 200 µL of blocking buffer in low-binding Eppendorf 1.5 mL tubes. We pelleted sorted nuclei at 1,000xg for 10 minutes at 4˚C, followed by resuspension in 50 µL Proteinase K Cleanup Buffer (recipe below) and 37˚C incubation for 30 minutes, and then freezing at -20˚C until library prep and sequencing.

For library prep, we purified tagmented DNA with 1.8x vol/vol Ampure XP beads (Beckman-Coulter catalog # A63881), eluted DNA in 11 µL and then PCR-amplified with Nextera Index kit primers (#FC-121-1012 from Illumina) using KAPA HiFi HotStart ReadyMix (KAPA Biosystems #KK2602) in a 30 µL reaction (72° 3:00, 95° 1:00, cycle 17x [98° :20, 65° :15, 72° :15], 72° 1:00). We purified PCR products using 1.8x Ampure XP beads, and quantified libraries using Agilent BioAnalyzer High Sensitivity DNA Chips (catalog # 5067-4626). Then sample libraries were pooled evenly and sequenced with paired-end 50bp reads either on Illumina MiSeq (Allen Institute) or NextSeq machines (SeqMatic, Fremont CA USA). We processed fastq files as described below.

*Single nuclear ATAC-seq.* We modified the single nuclear ATAC-seq workflow from the bulk sample workflow in several ways, most notably performing transposition reactions following sorting rather than prior to sorting, and omitting DAPI except for non-neuronal samples (due to the uncertainty of DAPI possibly interfering with transposition).

We collected and dissected specific MTG tissue layers as for bulk samples, but we immediately dounced the layers to release nuclei, and then stained in blocking buffer containing PE-NeuN antibody but not DAPI. We sorted single $NeuN^+$ nuclei from each layer into wells of a 96-well plate, using scatter profiles to exclude debris and doublets. We confirmed single nucleus-to-event correspondence by test-sorting single $NeuN^+$ events into flat-bottom 96 well plates with 40 µL blocking buffer containing DAPI followed by pelleting 1 min at 3,000xg and microscopic examination. These tests routinely yielded >95% single nucleus-filled wells and undetectable doublets. In the cases where glial cells were sorted, we first sorted neurons from the sample using PE-$NeuN^+$ staining, and then treated with DAPI (1 µg/µL) for 1-2 minutes prior to sorting glial cells as $DAPI^+NeuN^-$ events.

We sorted single $NeuN^+$ cells into 1.5 µL of Nextera Tn5 transposition reaction (0.6 µL Tn5 enzyme, 0.75 µL tagmentation buffer, 0.15 µL 1% IGEPAL CA-630) in Eppendorf semi-skirted 96-well plates (MilliporeSigma catalog # EP0030129504). Immediately following sorting we briefly centrifuged plates, vortexed, centrifuged plates again, and then incubated plates at 37˚C for 30 minutes for transposition. After transposition we added 0.6 µL Proteinase K Cleanup Buffer (recipe below), vortexed briefly and centrifuged, and incubated at 40˚C for an additional 30 minutes, then froze plates at -20 ˚C until library prep. Library prep for single nuclear samples was the same as for bulk samples, except we increased the number of amplification cycles from 17 to 22 cycles due to the lower input DNA content.

*Bulk ATAC-seq sample clustering.* We called peaks on all 39 bulk samples from five independent specimens using MACS2 (Zhang et al., 2008), and then used DiffBind (Ross-Innes et al., 2012) to identify 73,742 differential peaks for all contrasts among the sample types (sort strategies and specimens). Of these, 1,524 distinguished experimental specimens and were discarded for clustering. With 72,218 remaining peaks found specifically to discriminate any pairwise combinations of sort strategies, we reanalyzed correlation among bulk samples using reads in these peaks. This correlation matrix revealed groupings of non-neuronal samples, upper layer neuronal samples, and lower layer neuronal samples (Fig. S2C). One sample was omitted from this analysis (H17.03.009 L1 NeuN+) because this sample appeared intermediate between NeuN+ and NeuN- cells, likely due to a sorting error.

*ATAC-seq data preprocessing and quality control.* We retrieved sample-specific fastq files using standard built-in Illumina de-indexing protocols. We mapped each fastq file to human genome reference hg38 patch 7 using bowtie2 and the flags --no-mixed --no-discordant -X 2000 to generate sample-specific bam files, which we then filtered for low-quality mappings, secondary mappings, and unmapped reads using samtools view -q 10 -F 256 -F 4, and then filtered for duplicate reads using samtools rmdup. We then converted these filtered reads bam files to bed files using bedTools bamToBed for quality control calculations of mean ENCODE overlap and TSS enrichment score. For mean ENCODE overlap we converted bed files to fragment format, and assessed the percentage of unique fragments that overlap with ENCODE project DNaseI hypersensitivity peaks from adult human frontal cortex (studies ENCSR000EIK and ENCSR000EIY, (The ENCODE Project Consortium, 2012; Sloan et al., 2016) using bedTools intersectBed (Quinlan and Hall, 2010), and took the mean of these two numbers. For TSS enrichment score we used the published technique of Chen et al (Chen et al., 2016). This technique sums the overlap of reads in 2kb windows surrounding all human TSSs, then segments this 2kb window into 40 50-bp bins, then normalizes the summed read counts to the outside four bins (first and last two), and finally reports the TSS enrichment score as the maximum height of that normalized read count graph. We noticed that this technique worked well for all bulk samples but gave spurious abnormally high scores for some single nuclei having low read count; as a result we made the modification to set TSS enrichment score to 1 (no enrichment) for single nuclei having fewer than 500 reads or TSSs calculated to be greater than 20 (likely spurious events).

We used these quality control metrics to filter out low quality nuclei (ENCODE overlap < 15% AND TSS score < 4, Fig. S3). Additionally, we filtered out nuclei having fewer than 10,000 unique read pairs, since we require this many reads for our clustering approach. Of 3,660 initial cells we confined analysis to 2,858 high quality nuclei for clustering.

*Clustering single nuclei: bootstrapped clustering.* We clustered single nuclei using extended fragment Jaccard distance calculations among cells as implemented by the lowcat package (Graybuck et al., 2019). To accomplish this, we first excluded reads on chromosomes X, Y, and M to prevent differential chromosome-biased clustering. Then we randomly down-sampled to 10,000 unique fragments per nucleus, and then these fragments were extended to a regularized length of 1,000 bp with the same center. With these lists of extended fragments we next calculated the Jaccard similarity score for each nucleus pair, defined as the quotient of the intersecting extended fragment number, by the extended fragment union number. Then we calculated Jaccard distances among all nucleus pairs as 1 minus Jaccard similarity score.

Finally, this 2,858 x 2,858 Jaccard distance matrix was dimensionality reduced to a 2858 x 29 matrix of principal component variates, using axes 2 through 30 calculated by princomp in the R base stats package. We omitted principal component 1 because it was highly correlated to quality control metrics, suggesting that this axis primarily reflected library quality (Fig. S4B-D). Principal components beyond 30 contain little cell type information, so excluding them represents a de-noising step (Fig. S4A). These resulting 29 PCs are used to call nuclear clusters and to visualize them using tSNE.

To call cell clusters on this 2,858 x 29 principal component matrix, we bootstrapped an iterated PCA then Jaccard-Louvain clustering technique using k = 15 nearest neighbors (after testing k = 5,10,15,20, and finding 15 to give best visual separation of clusters on tSNE coordinates). We repeated each bootstrapping round 200 times, each time including only 80%

(2,286) of the nuclei, then performing PCA and using components 2 through 30 for Jaccard-Louvain clustering. Finally, we tabulated the frequency with which each nucleus co-clusters with every other nucleus. This co-clustering frequency matrix was then hierarchically clustered by Euclidean distances, and 27 cell type clusters were called by manually cutting the tree using idendr0 (https://github.com/tsieger/idendr0) to represent visually apparent co-clustered blocks of nuclei (Fig. S4E, left). Manual tree-cutting outperformed automatic tree cutting with cutree in the R stats package using either branch height or cluster number specified, likely since clusters have nonuniform separation and tightness.

Next we repeated this process with more stringent bootstrapping criteria: changing the percentage of cell to be re-clustered from 50-90%, and this analysis resulted in similar cluster structure and nucleus membership (Fig. S4E, middle, and Fig. S4F). In contrast, randomizing the Jaccard distance matrix prior to bootstrapped clustering yielded no clusters in the dataset (Fig. S4E, right). Together these analyses suggest that our identified clusters represent real and reproducible cell groups.

*Clustering single nuclei: comparing choice of feature set.* We also attempted to cluster nuclei using other feature sets besides Jaccard distances among cells (Fig. S4G). These additional feature sets included: 1) the list of all detected peaks from the entire aggregated dataset (236,588 peaks called using Homer findPeaks (Heinz et al., 2010) with -region flag), 2) the list of all RefSeq gene TSS regions, extended +/- 10kb (27,021 regions), 3) all 321,184 non-overlapping 10kb windows across the human genome, and 4) the list of "gene bins" defined as the genomic region for each gene between the boundaries of midpoints between each RefSeq gene transcribed region. For each feature set, we computed counts in features for each cell, then identified principal components, and visualized groupings by tSNE of principal components 2:50. For our dataset, Jaccard distances disclosed the qualitatively cleanest separation among nuclei, and among clusters (Fig. S4G). Furthermore, a wide range of tSNE perplexity values maintained these separations (Fig. S4H).

*Mapping clusters to transcriptomic cell types: assimilating epigenetic and transcriptomic information.* We wished to map our 2,858 high quality ATAC-seq profiled cells to human brain cell types discovered by large-scale RNA-seq studies (Hodge et al., 2019). To do this we first sought the best technique to manufacture gene-level information from the ATAC-seq data, in order to correlate with RNA-seq transcript counts. We tried four techniques: 1) read counts in RefSeq "gene bins" as above, 2) read counts in RefSeq gene bodies, 3) read counts in RefSeg gene TSS regions extended +/- 10 kb, and 4) Cicero gene activity scores (Cusanovich et al., 2018; Pliner et al., 2018). With these four sets of gene-level information computed for the 10000 fragment-downsampled library from each nucleus, we then mapped nuclei to RNA-seq cell types as the best correlated (highest Spearman correlation statistic) RNA-seq cluster (using median gene counts per million, CPM) with each nucleus, using each of four gene-level information vectors, resulting in four distinct mappings for each nucleus.

We calculated this correlation using a set of 831 marker genes, which we chose to be both informative marker genes for RNA-seq clustering and to contain abundant epigenetic information. This was accomplished by using the select_markers function with default parameters from the scrattch.hicat R package (Tasic et al., 2018) which yielded 2,791 transcriptomic marker genes, which was further filtered by intersecting with the top ten percent of genes with the highest summed Cicero gene activity scores across all 2,858 cells, to yield 831 combined transcriptomic and epigenetic marker genes for mapping.

The four sets of cellwise mappings yielded four tables of cell type abundances within our dataset. Next, taking the RNA-seq dataset (Hodge et al., 2019) as a true gold standard, we compared the four cell type abundance tables with the 'expected' cell type abundances, which was calculated as the sum of numbers of cells sorted in each sort strategy, times the expected cell type frequencies in each sort strategy. Correlating the four cell type abundance tables with the expected abundance table (pearson correlations of log-transformed abundance values plus one) revealed that, of the four techniques to compute gene-level information from ATAC-seq data, Cicero gene activity scores supply the most dependable gene-level information for the purpose of epigenetic to transcriptomic mapping (Fig. S5A).

*Mapping clusters to transcriptomic cell types: bootstrapping mapping for final mapping calls.* Using Cicero gene activity scores, we bootstrapped the cellwise mapping procedure 100 times with retention of a variable 50-90% of genes each round and applied the most frequently mapped transcriptomic cell type to each single ATAC-seq nucleus. Then we report the percentage of each cluster's constituent cells mapping to each cell type in Fig. S5B, and summed by cell type subclass in Fig. S5D.

We also performed clusterwise mapping for each of the 27 ATAC-seq clusters using the same bootstrapped mapping procedure, except that we aggregated Cicero gene activity scores by mean across cells within each cluster prior to mapping. We report the number of 100 times that each cluster is mapped to each cell type in Fig. S5C, and summed by transcriptomic subclass in Fig. S5E.

We observe that clusterwise mapping largely agrees with, but is cleaner than, cellwise mapping (compare Fig. S5B and S5C, also S5D and S5E, and S5F); hence we elect clusterwise mapping as the final mapping procedure. Each cell is thus assigned a final mapped cell type subclass (shown in Fig. S5E) as a result of its ATAC-seq cluster membership. For all downstream analyses of peaks, we use aggregations at the cell type subclass level as in Fig. S5E.

*Peak calling*. We called peaks on both bulk and aggregated single-nucleus data using Homer findPeaks with -region flag (Heinz et al., 2010). We found this program to be superior to Hotspot (v4), MACS2 (Zhang et al., 2008), and SICER (https://home.gwu.edu/~wpeng/Software.htm) to identify small regions corresponding to likely enhancers, while still capturing the peak boundaries. In preliminary experiments we observed that Hotspot returned small regions of a constant size (150bp or 250bp) that did not always align to peak summits, but it was relatively insensitive to read depth. MACS2 performed better than hotspot at picking full peak sizes but peak numbers found were strongly dependent on read depth. SICER returned very large regions (median >2kb) that did not clearly correspond to visual peaks. Using Homer findPeaks with -region flag, peak sizes are median 300-500 bp across subclasses, and we observed only a shallow dependence of identified peak number on read depth.

*Identifying transcription factor motifs using chromVAR.* We used chromVAR (Schep et al., 2017) to identify transcription factor motif accessibilities in our single nuclei. Using Homer findPeaks with -region flag, we called peaks on the aggregation of all single nuclear and bulk libraries (236,588 peaks), and then resized them to a standard 150bp size with the same center. We downloaded 452 transcription factor motifs from JASPAR (using JASPAR2018 R package, Tan, 2017), and 1,764 from cisBP (as included in the R package chromVARmotifs, Schep et al., 2017), and used chromVAR to aggregate and quantify motif accessibilities in all 2,858 single nuclei. Cell type subclass-distinguishing motifs across were found by ranking subclass-averaged motif accessibilities by standard deviation across subclasses (including DLX1, NEUROD6, RORB, and EMX2, Fig. S6A-D).

*Characterization of peaks by conservation.* With peaks called for each subclass, we calculated their phyloP scores as a measure of conservation. For peak phyloP scores, we used bigWigSummary to lookup phyloP values from hg38.phyloP4way.bw (Karolchik et al., 2004). These files quantify the basepair conservation across four mammals: *Homo sapiens*, *Mus musculus*, *Galeopterus variegatus* (Malayan flying lemur), and *Tupaia chinensis* (Chinese tree shrew). We return ten values evenly spaced across each peak, and calculate the maximum mean of eight three-consecutive-value sets. This is done to find smaller regions on the order of 100 bp highly conserved regions within each peak, and this technique yields greater deviations between real and random phyloP scores than taking a single peak-wise average alone. To compare conservation across groups of peaks, we subtracted the mean phyloP scores of randomized peak positions, from real peak phyloP scores (as in Fig. 2C).

*Identifying transcriptomic cell type matches for methylation data.* Using the dataset of Luo et al. (Luo et al., 2015), we correlated the published mCH gene body marker genes (their Supplementary Table 3 containing 1012 human and 1016 mouse methylation marker genes) with cluster-wise medians for transcriptomic human cell types (Hodge et al., 2019) and for mouse cell types (Tasic et al., 2018). We confined correlation analysis to the top 200 methylation marker genes published by Luo et al. that also have highest variance among transcriptomic cell subclasses. With these genes, we then calculated Pearson correlation coefficients between normalized gene body mCH and RNA-seq clusterwise median CPM, and assigned the best matches as the most anti-correlated mCH and CPM vectors as in Fig. S7. Specificity of matches was calculated as the difference between the best anti-correlation and the second-best anti-correlation. This analysis was repeated for both human and mouse datasets independently. Importantly, our transcriptomic cell type assignments agree with the previously predicted subclasses by Luo et al.

*Quantifying ATAC-seq peak overlaps with DMRs*. We first aggregated human DMRs from Luo et al. and Lister et al. (Lister et al., 2013; Luo et al., 2015). For neuron types, we downloaded DMRs and merged them using bedtools mergeBed. For non-neuron types, we downloaded raw fastq files from the GEO submission of Lister et al. (Lister et al., 2013) corresponding to bulk NeuN-negative cells from two human replicates (GSM1173774 and GSM1173777), and converted these to allc files using the pipeline analysis method of Luo et al. (Luo et al., 2017). These allc files were aggregated and used to find DMRs with methylpy DMRfind (minimum differentially methylated sites = 1) against allc files for all human subclasses from Luo et al., and an outgroup of human H1 cells from ENCODE. The same set of bulk non-neuronal DMRs were used for comparison to the ATAC-seq data for Astrocytes, Oligodendrocytes/OPCs, and Microglia subclasses (Fig. S6I-J).

With bed files corresponding to each subclass ATAC-seq peakset and to each subclass DMR set, we used bedtools intersectbed to quantify the overlap between peaks and DMRs. We bootstrapped calculation of real peak overlaps 100x by removing 20 percent of peaks each time and calculating percentage overlap, and the mean of these 100 measurements is reported. Similarly, we randomized peak positions throughout the genome 100x using bedtools shuffleBed, calculated percentage overlap each time, and the mean of these 100 measurements is reported. By definition, disjoint ranges of real versus randomized peak overlap percentages established false discovery rate < 0.01. We also calculated enrichment of DMR overlaps for ATAC-seq peaksets, defined as the ratio of real peak-DMR overlap percentage to the overlap percentage of randomized peak positions.

*Mouse to human cross-species comparisons.* We used the sets of subclass-specific (uniquely identified in only that subclass) peaks to map between human and mouse subclasses. We first mapped subclass-specific mouse peaks to hg38 using liftOver. Then we bootstrapped calculation of human peak overlap against all mouse peaks 100x with random retention of 80% of human peaks each time, and we took mean of Jaccard similarity coefficients (intersection over union) over 100 runs. In addition, we shuffled genomic peak positions 100x, and calculated mean Jaccard similarity coefficients each time. We report the enrichment of Jaccard similarity coefficients as the ratio of the real over random (Fig. 2A). To visualize set intersections in Venn diagram format we display results using all mouse and human peaks (not subclass-specific, Fig. 2B).

For characterization of human conserved and divergent peaks, we start with all human peaks and partition to those intersecting (“Conserved”) or not intersecting (“Divergent”) with mouse peaks identified within the same orthologous subclass and mapped to hg38 by liftOver. To characterize mouse conserved and divergent peaks, we intersect all mouse peaks with reciprocal mm10-mapped human peaks. Then we calculated phyloP scores as above.

*De novo sequence motif identification.* We used all mouse peaks and all human peaks to identify enriched sequence motifs using MEME-CHIP (Bailey et al., 2009). These motifs were then matched against known TF motifs in HOCOMOCO database v11 using TomTom. We then filtered the MEME-CHIP output by first excluding all motifs with $-\log_{10}$(E-value) < 5; E-value represents the enrichment p-value (by Fisher's exact test) times the number of candidate motifs tested. We further filtered by second excluding all motif matches with TFs not expressed (median CPM = 0) in that cell subclass from RNA-seq studies (Tasic et al., 2018; Hodge et al., 2019). Third we filtered by excluding all low-confidence motif matches with E-value > 0.2 and q-value > 0.2; q-value represents the minimal false discovery rate at which the observed similarity would be deemed significant. Finally, these filtered lists of all detected motifs in all cell subclasses were manually curated to a master list of all high-confidence identified TF motifs.

*Quantifying repetitive element overlap.* To characterize the repetitive element overlap for peaks, we first partitioned mouse and human subclass-specific peaksets to conserved and divergent peaks. Then we calculated the overlap with repetitive genomic elements using hg38 and mm10 RepeatMasker (v.4.0.5, http://www.repeatmasker.org) files, using a 100x bootstrapped overlap and 100x bootstrapped randomization strategy as described above for DMR overlap. Human L56IT peaks were omitted from this analysis because very few of these peaks are both subclass-specific and conserved.

*Cloning enhancers.* Enhancers were chosen for cloning from epigenetic data using one of two strategies. For the first strategy we used the following criteria: 1) visible specific peak manually identified in read pileups adjacent to known subclass-specific marker genes, and 2) containing a region of high primary sequence conservation by phyloP score. For the second strategy we used the following criteria: 1) a subclass-specific ATAC-seq peak identified by Homer (with -region flag) in both human and mouse (conserved) or only human (divergent), 2) a subclass-specific DMR in both human and mouse (conserved) or only human (divergent), 3) ranking by human ATAC-seq read counts within region, and 4) manual confirmation by visualization of read pileup by experimenter.

Chosen enhancers were cloned into either scAAV or rAAV (ssAAV) expression vectors. For scAAV vectors we used a plasmid backbone that is a derivative of pscAAV-MCS (Cell

Biolabs catalog # VPK-430, for scAAV vectors), which was used for eHGT_017h, eHGT_019h, eHGT_022h/m, eHGT_023h, eHGT_025h, and hDLXI56i (Dimidschstein et al., 2016; Chan et al., 2017). For rAAV (ssAAV) vectors we used a plasmid backbone from Addgene plasmid number 51084 (AAV-hSyn1-GCaMP6s-P2A-nls-dTomato, which was itself originally derived from pAAV-GFP [Cell Biolabs catalog # VPK-410]). We used this rAAV backbone for eHGT_058h, eHGT_064h, eHGT_078h/m, eHGT_079h, eHGT_082h, eHGT_096h, eHGT_098h, eHGT_128h, and eHGT_140h, hDLXI56i, and 3x(hDLXI56i core). Enhancers were inserted by standard Gibson assembly approaches, upstream of a minimal beta-globin promoter and the reporter SYFP2, a brighter EGFP alternative that is well tolerated in neurons (Kremers et al., 2006). NEB Stable cells (New England Biolabs # C3040I) were used for transformations and cultured at 32°C. scAAV plasmids were monitored by restriction analysis and Sanger sequencing for occasional recombination of the left ITR; this left ITR recombination was not observed for rAAV plasmids. We attempted to boost expression level for some enhancers by engineering a triple tandem array of the enhancer core ("concatemer"), for example for 3x(hDLXI56i core) as in Figure 5H,K.

*Virus production*. Enhancer AAV plasmids were maxi-prepped and transfected with PEI Max 40K (Polysciences Inc., catalog # 24765-1) into one 15 cm plate of AAV-293 cells (Cell Biolabs catalog # AAV-100), along with helper plasmid pHelper (Cell BioLabs) and PHP.eB rep/cap packaging plasmid (Chan et al., 2017), with a total mass of 150 µg PEI Max 40K, 30 µg pHelper, 15 µg rep/cap plasmid, and 15 µg enhancer-AAV vector. The next day medium was changed to 1% FBS, and then after 5 days cells and supernatant were harvested and AAV particles released by three freeze-thaw cycles. Lysate was then treated with benzonase to degrade free DNA (2 µL benzonase, 30 min at 37°C, MilliporeSigma catalog # E8263-25KU), and then cell debris was cleared with low-speed spin (1500g 10 min). The supernatant containing virus was concentrated over a 100 kDa molecular weight cutoff Centricon column

(MilliporeSigma catalog # Z648043) to a final volume of ~150 µL. For highly purified large-scale preps this protocol was altered so that ten plates were transfected and harvested together at 3 days after transfection, and then the crude virus was purified by iodixanol gradient centrifugation.

*Mouse virus testing*. Mice were retro-orbitally injected at P42-P49 with 10 µL (approximately $1 \times 10^{11}$ genome copies) of crude virus prep diluted with 100 µL PBS, then sacrificed at 18-28 days post infection. For live epifluorescence, we perfused mice with ACSF.7 and cut live 350 µm sections with a compresstome from one hemisphere to analyze reporter expression using a 10x objective on a Nikon Ti-Eclipse epifluorescence microscope with built-in real-time deconvolution image processing for thick tissues (Nikon Image Systems Elements software with Advanced Research module). For full sagittal section images of mouse brain, we processed the brain for mFISH HCR and anti-GFP immunostaining (as below), and using a 4x objective on an Olympus FV3000 confocal we took images in a 3x5 grid tiling the brain at two optical slices separated by 4 µm, and in ImageJ we performed z-projections using maximum intensity and stitched images using Grid stitching and linear blending fusion method. For antibody staining the other hemisphere was drop-fixed in 4% PFA in PBS for 4-6 hours at 4°C, then cryoprotected in 30% sucrose in PBS 48-72 hours, then embedded in OCT for 3 hours at room temperature, then frozen on dry ice and sectioned at 10 µm thickness, prior to antibody stain using standard practice. We used the following primary antibodies: chicken anti-GFP (Aves # GFP-1020), rabbit anti-Parvalbumin (Swant # PV27), rabbit anti-Somatostatin (Peninsula Biolabs # T-4547), rabbit anti-VIP (BosterBio # RP1108), and mouse anti-RFP (abcam # ab65856) to detect mCherry from *Gad2-T2a-NLS-mCherry* mice (Peron et al., 2015). Secondary antibodies were 488-, 555-, and 647-conjugated secondary antibodies from ThermoFisher Scientific. We performed single-cell RNA-seq from the mouse visual cortex as described previously (Tasic et al., 2016, 2018).

*Multiplexed FISH by hybridization chain reaction (mFISH-HCR)*. We performed this technique on mouse brain hemispheres fixed by immersion in 4% PFA in PBS for 4-6 hours at 4°C. After fixation, we rinsed hemispheres with PBS and stored them in PBS at 4°C for up to one month. For sectioning, we embedded hemispheres in 1% low-melt agarose in PBS and cut 50 µm sagittal sections on a Leica VT1000S vibratome in cold PBS buffer. We post-fixed sections in 4% PFA in PBS for 2 hours and then rinsed in PBS at room temperature, then dehydrated with 70% ethanol at 4°C. Afterwards sections could be stored for up to a month in 4°C. For staining, we cleared sections with 8% SDS in PBS for 2 hours at room temperature then washed three times in 2x SSC for 1 hour each, then with Hybridization Buffer (Molecular Instruments) in a new well before applying Hybridization Buffer containing HCR Probes and hybridized overnight at 37°C. The next day we washed samples with 30% Probe Wash Buffer for 1 hour at 37°C, then rinsed with 2xSSC. During the probe wash, we denatured fluorescently labeled HCR hairpins at 95°C for 90 seconds and then snap-cooled in a room temperature aluminum block tube holder for 30 minutes. We added the denatured hairpins to Amplification Buffer and applied to tissue sections for 2 hours at room temperature in the dark, then washed with 2x SSC containing DAPI, again with 2x SSC, and finally mounted on SuperFrost Plus slides in Prolong Glass Mounting medium (Thermo Fisher Scientific # P36980). We imaged these HCR stains with an Olympus FV3000 confocal microscope using manufacturer's software. Molecular Instruments generated HCR probes against the following transcripts:  Rorb NM_001043354.2; Lamp5 NM_029530.2; Vip NM_011702.3; Pvalb NM_001330686.1; Sst NM_009215.1; Slc17a7 NM_182993.2; Gad1 NM_008077.5.

*Human ex vivo AAV vector testing.* We transported neurosurgical temporal cortex samples from the operating suite to the Allen Institute in typically less than 30 minutes, using specialized transportation equipment to maintain sterility and carbogen bubbling throughout processing. Tissue samples were blocked and then sliced at 350 µm thickness and white matter and pial membranes were dissected away. Slices then underwent warm recovery (bubbled ACSF.7 at 30 degrees for 15 minutes) followed by reintroduction of sodium (bubbled ACSF.8 at room temperature for 30 minutes, recipe below, (Ting et al., 2018). Slices were then plated at the gas interface on Millicell PTFE cell culture inserts (MilliporeSigma # PICM03050) in a 6-well dish on 1 mL of Slice Culture Medium (recipe below). After 30 minutes, slices were infected by direct application of high-titer AAV2/PHP.eB viral prep to the surface of the slice, 1 µL per slice. Slice culture medium was replenished every 2 days and reporter expression was monitored. For eHGT_022h in human, imaging and single cell RNA-seq was performed at 42 days *in vitro*. For hDLXI56i in human, imaging and single cell RNA-seq was performed in four independent experiments at 8, 13, 28, and 69 days *in vitro.* A fifth experiment on hDLXI56i (at 11 days *in vitro*) was excluded from analysis because 36/48 (75%) of sorted cells either failed to map to transcriptomic cell types, or mapped as uncertain non-neuronal types; this is likely due to either a failed sort or poor starting tissue quality given heterogeneity of patient samples.

Single cell RNA-seq was accomplished on human virus-infected neurons by 1 hour digestion at 30ºC in carbogenated ACSF.1/trehalose + blockers + papain (all recipes below), followed by gentle trituration in Low-BSA Quench buffer, shallow spin gradient centrifugation (100g 10 minutes at room temperature) into High-BSA Quench buffer, and resuspension into Cell Resuspension Buffer. We also employed Myelin Bead Removal Kit II (Miltenyi catalog # 130-096-733) at 1/20 to remove myelin debris, and PE-anti CD9 clone eBioSN4 (Thermo Fisher catalog # 12-0098-42) at 1/50 to sort away contaminating glial cells. Then we sorted single SYFP2$^+$ labeled human neurons for sequencing using SMARTer V4 as previously described (Tasic et al., 2016, 2018). To map single cells to the transcriptomic taxonomies, we trained a nearest centroid classifier on cell type labels using human and mouse V1 scRNA-seq cluster labels (Tasic et al., 2018), employing informative marker genes chosen by the select.markers function in scrattch.hicat (Tasic et al., 2018). We confined taxonomy mapping analysis to the cells that passed cDNA library generation quality control metrics and showed detectable levels of SYFP2 transcripts. Intermediate-mapping cells are represented as circles on nodes of the cluster dendrograms.

*In vivo non-human primate AAV vector testing.* All procedures used with macaque monkeys conformed to the guidelines provided by the US National Institutes of Health and were approved by the University of Washington Animal Care and Use Committee. One rhesus macaque (*Macaca mulatta*) was injected with an AAV vector in ten injection sites during a single surgery. Two sites were located in left temporal cortex, three sites each in left and right occipital cortex, and one site each in left and right motor cortex. AAVs were purified by iodixanol gradient ultracentrifugation for this procedure. After craniotomy, using a pneumatic pico pump (World Precision Instruments) a total of 5 µL AAV vector was injected at each site with 500 nL expelled at each of ten depths evenly spaced from 2 mm to 200 µm deep beneath the pial surface. Sites were separated by ~1 cm in each region with multiple injection sites. Four sites are described in this manuscript (eHGT_079h, 128h, and 140h in occipital cortex, and 140h in temporal cortex). Fifty days after injection, the animal was sacrificed. We inspected the brain surface, cut tissue blocks (~2x2x2cm) around each visible fluorescent spot, and fixed each block 4% PFA in PBS for 24 hours at 4ºC. After PFA fixation, we embedded blocks in 2% agarose in PBS and cut 350 µm sections and inspect each for fluorescent cells. We then cryopreserved a subset of sections in 30% sucrose in water overnight and subsectioned them on a sliding microtome to 30 µm for immunostaining using the following antibodies: chicken anti-GFP (Aves # GFP-1020), rabbit anti-Parvalbumin (Swant # PV27), and guinea pig anti-GABA (EMD Millipore # AB175). Images shown are from the region of high labeling close to the needle tract (<1 mm), but the zone of expression extended for ~3-4 mm orthogonal to the needle tract. Proper recovery of sites was confirmed by PCR on DNA from dissected fixed thick slices (recovered with QIAamp DNA FFPE Tissue Kit, Qiagen catalog # 56404 ) using common primers to all vectors: F 5'-ACTCCATCACTAGGGGTTCCTG and R 5'-GGACACGCTGAACTTGTGGC followed by Sanger sequencing with the nested reverse primer 5'-ACGTCGCCGTCCAGCTC.

*Ex vivo non-human primate AAV vector testing.* Brains from healthy *Macaca nemestrina* animals housed at the Washington National Primate Research Center (Seattle, WA) aged 2-15 years were obtained through the Tissue Distribution Program. Whole hemispheres or tissue blocks were transported to the Allen Institute and processed for *ex vivo* culture and AAV vector testing as described above for human neurosurgical samples (Ting et al., 2018). Data are shown for cultures of MTG tissue. Cell subclass selectivity was evaluated by mFISH-HCR as described above for mouse, except that 350 µm cultured slices were cleared with 67% 2,2′-thiodiethanol in water prior to mounting on slides for microscopy. Molecular Instruments designed probes with the following accession numbers provided to them: Slc17a7 NM_ 005589901.2; Gad1 NM_ 005573441.2; Vip NM_ 005552161.2; Pvalb NM_ 005567398.2; Sst NM_ 005545442.2.

*Inferring GWAS-cell subclass associations.* We used linkage disequilibrium score regression (LDSC, (Bulik-Sullivan et al., 2015; Finucane et al., 2015) to partition heritability of various brain conditions to regions associated with accessible chromatin in eleven human cortical cell subclasses, whose peaks are grouped into Conserved and Divergent subsets. As outgroup comparators, we also assessed the heritability associated with outgroup populations of human keratinocytes downloaded from ENCODE (The ENCODE Project Consortium, 2012). Additionally, we also performed this analysis using DMRs from human cortical neuron subclasses (Luo et al., 2017), human cortical non-neurons (Lister et al., 2013), and H1 human embryonic stem cells (The ENCODE Project Consortium, 2012).

Summary statistics from 21 GWAS studies were acquired and evaluated including brain-related (schizophrenia, major depressive disorder, autism spectrum disorder, ADHD, Alzheimer's disease, Tourette's syndrome, bipolar disorder, eating disorder, obsessive-compulsive disorder, loneliness, BMI, PTSD) and non-brain-related diseases (Crohn's disease and asthma) from the PGC and EMBL/EBI GWAS repositories (Anney et al., 2017; Autism Spectrum Disorder Working Group of the Psychiatry Genomics Consortium, 2015; Demenais et al., 2018; Demontis; Duncan et al., 2017, 2018; Gao et al., 2017; International Obsessive Compulsive Disorder Foundation Genetics Collaborative (IOCDF-GC) and OCD Collaborative Genetics Association Studies (OCGAS), 2018; Lambert et al., 2013; de Lange et al., 2017; Lee et al., 2018; Liu et al., 2015; Major Depressive Disorder Working Group of the Psychiatric GWAS Consortium et al., 2013; Marioni et al., 2018; Okbay et al., 2016; Psychiatric GWAS Consortium Bipolar Disorder Working Group, 2011; Schizophrenia Psychiatric Genome-Wide Association Study (GWAS) Consortium, 2011; Schizophrenia Working Group of the Psychiatric Genomics Consortium, 2014; Tourette Association of America International Consortium for Genetics (TAAICG, 2018; Wray et al., 2018; Yang et al., 2017).

We excluded studies with $\log_{10}( N * h^2 ) < 3.6$, where N is number of patients in the study and $h^2$ represents the sum of heritability across SNPs within the study, which represents the effective power of the study (Finucane et al., 2015). This exclusion removed 6 studies: asthma (Demenais et al., 2018), $\log_{10}( N * h^2 ) = 3.5$), PTSD (Duncan et al., 2018), $\log_{10}( N * h^2 ) = 2.9$), eating disorder (Duncan et al., 2017), $\log_{10}( N * h^2 ) = 3.5$), loneliness (Gao et al., 2017), $\log_{10}( N * h^2 ) = 3.3$), obsessive-compulsive disorder (International Obsessive Compulsive Disorder Foundation Genetics Collaborative (IOCDF-GC) and OCD Collaborative Genetics Association Studies (OCGAS), 2018), $\log_{10}( N * h^2 ) = 3.5$), and one major depressive disorder study (Major Depressive Disorder Working Group of the Psychiatric GWAS Consortium et al., 2013), $\log_{10}( N * h^2 ) = 3.3$). The 15 studies with sufficient power for inclusion were all performed on a European descent population. Within these datasets, we confined analysis to 1,389,227 high-confidence SNPs present in the HapMap3 list, and using linkage disequilibrium maps from the 1000 Genomes Project European descent individuals, we analyzed the trait and disease enrichments of cell subclass-associated chromatin along with the LDSC baseline model LDv2.0 with 75 enumerated genomic feature categories. LDSC was performed to associate these 15 studies with both ATAC-seq peaks and methylation DMRs (Fig. S8, Lister et al., 2013; Luo et al., 2017), and both epigenetic data modalities gave qualitatively similar results although ATAC-seq peaks give stronger enrichments. Generally weak associations were observed between the outgroup disease (Crohn's disease) with brain cell types, and between the outgroup peak set (Keratinocytes, Fig. S8A,C, The ENCODE Project Consortium, 2012) and brain diseases. For statistical testing of enrichments, we use Bonferroni multiple hypothesis testing correction of LDSC's block jackknife-estimated p-values, as previously suggested (Skene et al., 2018). This correction is 0.05 / 345 disease/subclass combinations = 1.45e-4 significance cutoff in Fig. S8C. We similarly use 180 and 150 tests in Fig. S8A and S8B.

*Buffer Recipes.*

<u>Proteinase K Cleanup Buffer</u>

| | |
|---|---|
| EDTA | 50 mM |
| Sodium chloride | 5 mM |
| Sodium dodecyl sulfate | 1.25% (w/v) |
| Proteinase K (Qiagen # 19131) | 5 mg/mL |
| pH 8.0 | |

<u>Nuclei Isolation Medium</u>

| | |
|---|---|
| Sucrose | 250 mM |
| Potassium chloride | 25 mM |
| Magnesium chloride | 5 mM |
| Tris-HCl | 10 mM |
| pH 8.0 | |

<u>Homogenization Buffer</u>

10 mL Nuclei Isolation Medium
0.1% (w/v) Triton X-100
One pellet Roche Mini cOmplete EDTA-free (Sigma catalog # 4693159001)

<u>Blocking Buffer</u>

PBS
0.5% (w/v) BSA (catalog # A2058 from Millipore Sigma)
0.1% (w/v) Triton X-100

<u>ACSF.7</u>

| | |
|---|---|
| HEPES | 20 mM |
| Sodium Pyruvate | 3 mM |
| Taurine | 10 µM |

| | |
|---|---|
| Thiourea | 2 mM |
| D-(+)-glucose | 25 mM |
| Myo-inositol | 3 mM |
| Sodium bicarbonate | 30 mM |
| Calcium chloride dihydrate | 0.5 mM |
| Magnesium sulfate | 10 mM |
| Potassium chloride | 2.5 mM |
| Monosodium Phosphate | 1.25 mM |
| HCl | 92 mM |
| N-methyl-D-(+)-glucamine | 92 mM |
| L-ascorbic acid | 5.0 mM |
| N-acetyl-L-cysteine | 12 mM |

pH adjusted to 7.3-7.4, osmolarity adjusted to 295-305, and carbogenated.

<u>ACSF.8</u>

| | |
|---|---|
| HEPES | 20 mM |
| Sodium Pyruvate | 3 mM |
| Taurine | 10 µM |
| Thiourea | 2 mM |
| D-(+)-glucose | 25 mM |
| Myo-inositol | 3 mM |
| Sodium bicarbonate | 30 mM |
| Calcium chloride dihydrate | 2.0 mM |
| Magnesium sulfate | 2.0 mM |
| Potassium chloride | 2.5 mM |
| Monosodium Phosphate | 1.25 mM |
| Sodium chloride | 92 mM |
| L-ascorbic acid | 5.0 mM |
| N-acetyl-L-cysteine | 12 mM |

pH adjusted to 7.3-7.4, osmolarity adjusted to 295-305, and carbogenated.

<u>Slice Culture Medium</u>

| | | |
|---|---|---|
| MEM Eagle medium powder | 1680 mg | (MilliporeSigma catalog # M4642) |
| L-ascorbic acid powder | 36 mg | |
| CaCl2, 2.0 M | 100 µL | |
| MgSO4, 2.0 M | 200 µL | |
| HEPES, 1.0 M | 6.0 mL | |
| Sodium bicarbonate, 893 mM | 3.36 mL | |
| D-(+)-glucose, 1.11 M | 2.25 mL | |
| Pen/Strep 100x (5k U/mL) | 1.0 mL | (Thermo catalog # 15070063) |
| Tris base, 1.0 M | 260 µL | |
| GlutaMAX 200 mM | 0.5 mL | (Thermo catalog # 35050061) |
| Bovine Pancreas Insulin, 10 mg/mL | 20 µL | (MilliporeSigma catalog # I0516) |
| Heat-inactivated horse serum | 40 mL | (Thermo catalog # 26050088) |

| | |
|---|---|
| Deionized water | to 250 mL |

pH adjusted to 7.3-7.4, and osmolarity adjusted to 300-305,

ACSF.1/trehalose

| | |
|---|---|
| HEPES | 20 mM |
| Sodium Pyruvate | 3 mM |
| Taurine | 10 μM |
| Thiourea | 2 mM |
| D-(+)-glucose | 25 mM |
| Myo-inositol | 3 mM |
| Sodium bicarbonate | 25 mM |
| Calcium chloride dihydrate | 0.5 mM |
| Magnesium sulfate | 10 mM |
| Potassium chloride | 2.5 mM |
| Monosodium Phosphate | 1.25 mM |
| Trehalose dihydrate | 132 mM |
| HCl | 2.9 mM |
| N-methyl-D-(+)-glucamine | 30 mM |
| L-ascorbic acid | 5.0 mM |
| N-acetyl-L-cysteine | 12 mM |

pH adjusted to 7.3-7.4, and osmolarity adjusted to 295-305.

ACSF.1/trehalose + blockers:
50 mL ACSF.1/trehalose
50 μL 100 μM TTX (final 0.1 μM)
100 μL 25 mM DL-AP5 (final 50 μM)
15 μL 60 mM DNQX (final 20 μM)
5 μL 100 mM (+)-MK801 (final 10 μM)

ACSF.1/trehalose + blockers + papain:
15 mL ACSF.1/trehalose + blockers
One vial Worthington PAP2 reagent (150 U, final 10U/mL)
15 μL 10kU/mL DNase I (Roche)

Low-BSA Quench buffer
15 mL ACSF.1/trehalose + blockers
15 μL 10kU/mL DNase I (Roche)
150 μL 20% BSA dissolved in water (final conc 2 mg/mL)
150 μL 10 mg/mL ovomucoid inhibitor (Sigma T9253, final concentration 0.1 mg/mL)

High-BSA Quench buffer
15 mL ACSF.1/trehalose + blockers
15 μL 10kU/mL DNase I (Roche)
750 μL 20% BSA dissolved in water (final concentration 10 mg/mL)

150 μL 10 mg/mL ovomucoid inhibitor (Sigma T9253, final concentration 0.1 mg/mL)

ACSF.1/trehalose + EDTA

| | |
|---|---|
| HEPES | 20 mM |
| Sodium Pyruvate | 3 mM |
| Taurine | 10 μM |
| Thiourea | 2 mM |
| D-(+)-glucose | 25 mM |
| Myo-inositol | 3 mM |
| Sodium bicarbonate | 25 mM |
| Potasium chloride | 2.5 mM |
| Monosodium Phosphate | 1.25 mM |
| Trehalose | 132 mM |
| HCl | 2.9 mM |
| EDTA | 0.25 mM |
| N-methyl-D-(+)-glucamine | 30 mM |
| L-ascorbic acid | 5.0 mM |
| N-acetyl-L-cysteine | 12 mM |

pH adjusted to 7.3-7.4, and osmolarity adjusted to 295-305.

Cell Resuspension Buffer

50 mL ACSF.1/trehalose + EDTA
50 μL 100 μM TTX (final concentration 0.1 μM)
100 μL 25 mM DL-AP5 (final concentration 50 μM)
15 μL 60 mM DNQX (final concentration 20 μM)
5 μL 100 mM (+)-MK801 (final concentration 10 μM)
150 μL 20% BSA dissolved in water (final concentration 2 mg/mL)
1 μg/mL 4'-diamino-phenylindazole (DAPI)

**A**

| Specimen | Experiments | Age | Gender | Disease | Region |
|---|---|---|---|---|---|
| H17.03.008 | Bulk | 60 | F | Epilepsy | MTG TCx |
| H17.03.009 | Bulk | 52 | M | Epilepsy | MTG TCx |
| H17.03.010 | Bulk | 38 | F | Epilepsy | MTG TCx |
| H17.26.001 | Bulk, single | 58 | F | Tumor | TCx |
| H17.26.003 | Bulk, single | 25 | M | Tumor | TCx |
| H17.06.012 | Single | 23 | M | Epilepsy | MTG TCx |
| H17.26.002 | Single | 57 | F | Tumor | TCx |
| H17.06.015 | Single | 19 | M | Epilepsy | MTG TCx |
| H18.03.001 | Single | 38 | F | Epilepsy | TCx |
| H18.03.005 | Single | 49 | F | Epilepsy | MTG TCx |
| H18.06.006 | Single | 38 | M | Epilepsy | TCx |
| H18.03.006 | Single | 24 | F | Epilepsy | MTG TCx |
| H18.03.008 | Single | 19 | F | Epilepsy | TCx |
| H18.03.010 | Single | 31 | M | Epilepsy | MTG TCx |
| H18.03.012 | Single | 22 | M | Epilepsy | MTG TCx |
| H18.26.405 | Single | 68 | F | Tumor | TCx |
| H18.03.313 | Single | 23 | F | Epilepsy | MTG TCx |

**B**

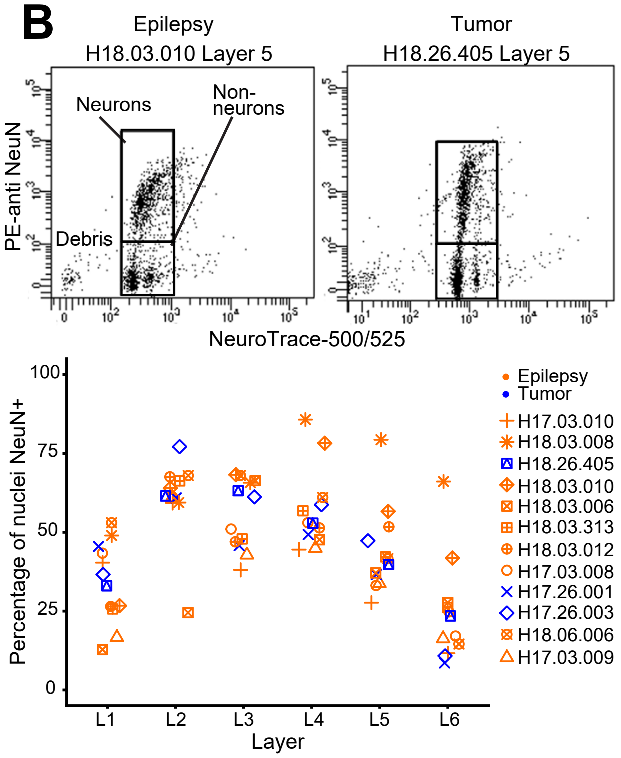

**Supplementary Figure 1: Profiling chromatin accessibility across multiple human temporal cortex tissue samples.**

A) Summary table of 17 human neurosurgical specimens used for chromatin accessibility profiling experiments by ATAC-seq. De-identified specimen codes are given along with type of ATAC-seq experiments performed (Bulk or Single-nucleus), age, gender, patient disease requiring surgery, and region of tissue harvested.

B) Flow cytometry analysis of sorted nuclei demonstrating that tumor and epilepsy cases display qualitatively similar-staining nuclei, and quantitatively similar proportions of neuronal nuclei. On top, example flow plots from PE-anti NeuN and NeuroTrace 500/525-stained layer 5 nuclei from one epilepsy and one tumor case. On bottom, percentages of nuclei labeled with anti-NeuN antibody from six dissected layers of cortex, from 12 specimens. Four specimens were not analyzed in this way, and one specimen (H18.03.005) was omitted from this analysis because of poor staining signal.

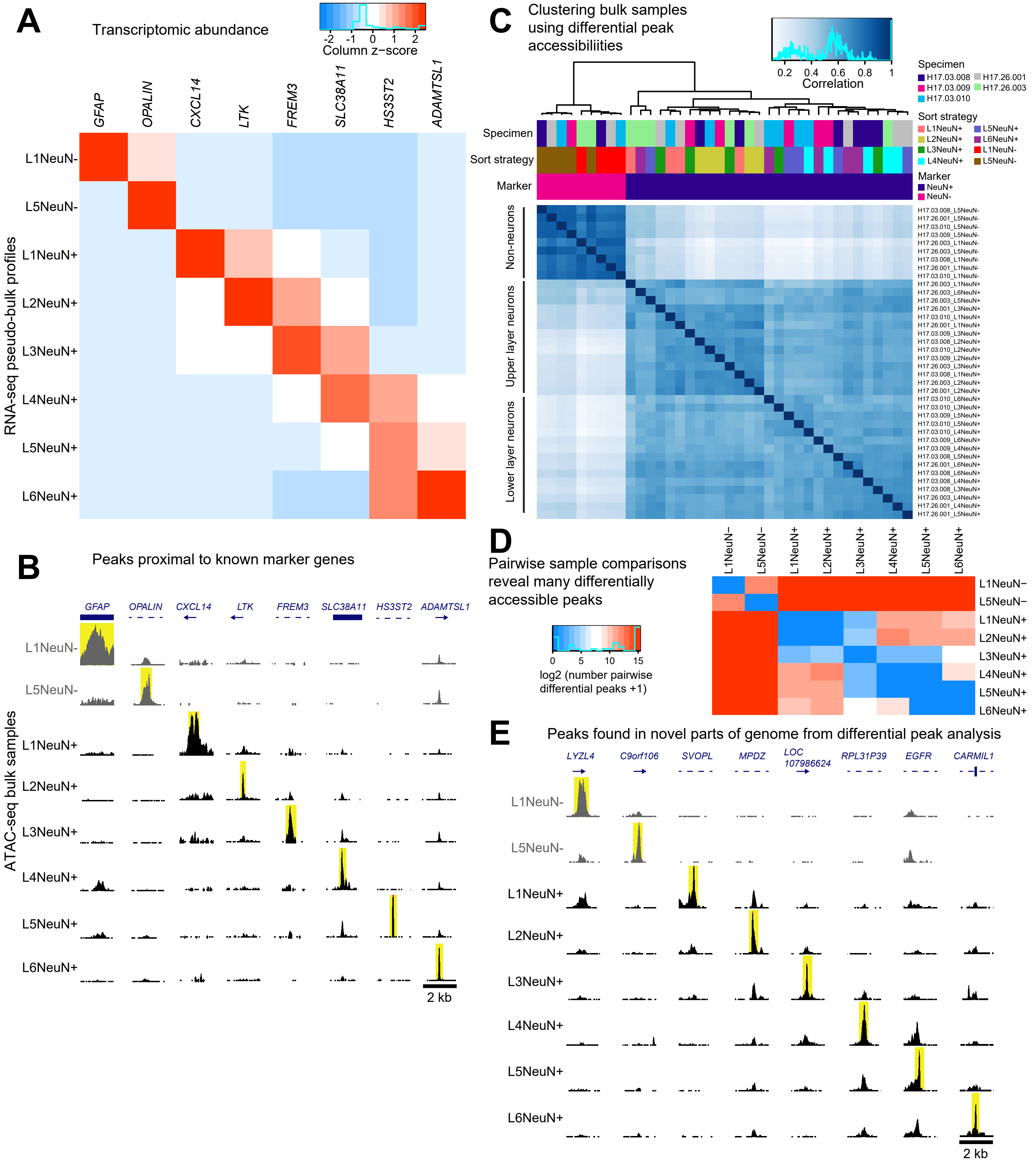
A
Transcriptomic abundance
-2 -1 0 1 2
Column z−score
GFAP
OPALIN
CXCL14
LTK
FREM3
SLC38A11
HS3ST2
ADAMTSL1
RNA-seq pseudo-bulk profiles
L1NeuN-
L5NeuN-
L1NeuN+
L2NeuN+
L3NeuN+
L4NeuN+
L5NeuN+
L6NeuN+
B
Peaks proximal to known marker genes
ATAC-seq bulk samples
2 kb
C
Clustering bulk samples using differential peak accessibiliities
0.2 0.4 0.6 0.8 1
Correlation
Specimen
H17.03.008
H17.03.009
H17.03.010
H17.26.001
H17.26.003
Sort strategy
L1NeuN+
L2NeuN+
L3NeuN+
L4NeuN+
L5NeuN+
L6NeuN+
L1NeuN-
L5NeuN-
Marker
NeuN+
NeuN-
Non-neurons
Upper layer neurons
Lower layer neurons
D
Pairwise sample comparisons reveal many differentially accessible peaks
0 5 10 15
log2 (number pairwise differential peaks +1)
L1NeuN−
L5NeuN−
E
Peaks found in novel parts of genome from differential peak analysis
LYZL4
C9orf106
SVOPL
MPDZ
LOC 107986624
RPL31P39
EGFR
CARMIL1
2 kb

**Supplementary Figure 2: Bulk ATAC-seq data demonstrates differentially accessible chromatin elements around known marker genes, and in novel genomic regions.**

A) snRNA-seq data (Hodge et al., 2019), aggregated into pseudo-bulk profiles by weighted averages of gene CPM medians for 75 transcriptomic clusters. Weights were assigned by their frequencies within the eight sort strategies, and the heatmap is scaled by z-score within each column (gene). Relative expressions of eight sort strategy-specific marker genes are displayed.

B) Example sort strategy-specific peaks proximal to (<50kb distance to gene body) the eight sort strategy-specific transcriptomic marker genes. Pileups indicate aggregated data within a 2 kb genomic window across five independent experiments. In B and E, dashed lines indicate introns, thick lines indicate exons, and arrows indicate direction towards proximal marker gene. Yellow highlights demarcate sort strategy-specific chromatin accessibility peaks.

C) DiffBind (Ross-Innes et al., 2012) identification of 72,218 peaks that were differentially accessible among any pairwise comparison of sort strategies (FDR 0.01). Read counts within those 72,218 differentially accessible peaks then clustered samples using a correlation distance matrix, which revealed separate groupings of non-neuronal samples, and upper- and lower-layer neuronal samples. One sample was omitted from this analysis (H17.03.009 L1 NeuN+) because this sample appeared intermediate between NeuN+ and NeuN- cells, suggesting a failed sort.

D) Number of peaks differentiating each pairwise sample contrast.

E) Example sort strategy-specific peaks resulting from pairwise DiffBind differential peak analysis. These peaks were found in novel genomic regions (not proximal to known marker genes), and closest genes are shown.

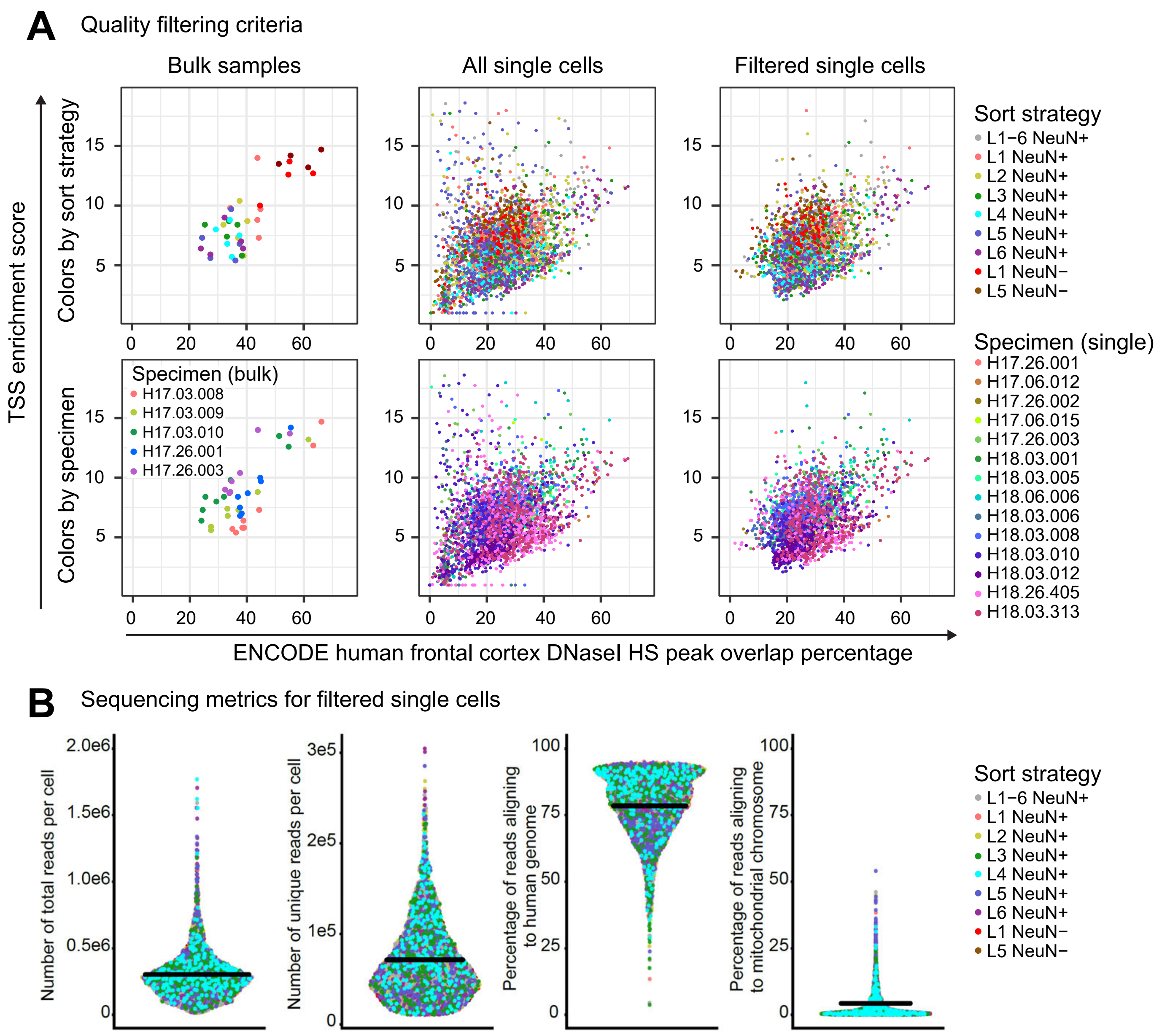
A
Quality filtering criteria
Bulk samples
All single cells
Filtered single cells
TSS enrichment score
Colors by sort strategy
Colors by specimen
Specimen (bulk)
H17.03.008
H17.03.009
H17.03.010
H17.26.001
H17.26.003
Sort strategy
L1−6 NeuN+
L1 NeuN+
L2 NeuN+
L3 NeuN+
L4 NeuN+
L5 NeuN+
L6 NeuN+
L1 NeuN−
L5 NeuN−
Specimen (single)
H17.26.001
H17.06.012
H17.26.002
H17.06.015
H17.26.003
H18.03.001
H18.03.005
H18.06.006
H18.03.006
H18.03.008
H18.03.010
H18.03.012
H18.26.405
H18.03.313
ENCODE human frontal cortex DNaseI HS peak overlap percentage
B
Sequencing metrics for filtered single cells
Number of total reads per cell
Number of unique reads per cell
Percentage of reads aligning to human genome
Percentage of reads aligning to mitochondrial chromosome
Sort strategy
L1−6 NeuN+
L1 NeuN+
L2 NeuN+
L3 NeuN+
L4 NeuN+
L5 NeuN+
L6 NeuN+
L1 NeuN−
L5 NeuN−

**Supplementary Figure 3: High quality single cell ATAC-seq libraries.**

A) Quality control metrics for bulk (left), all 3660 single nuclei (middle), and 2858 quality-filtered single nuclei (right). Metrics were ENCODE frontal cortex DNaseI hypersensitivity peak overlap percentage (x axis), and TSS enrichment score (y axis). Poor-quality single nuclei with TSS enrichment score < 4 AND ENCODE overlap < 15%, OR < 10000 unique mapped reads per nucleus were omitted. Plots are colored by sort strategy (top) or by specimen (bottom).

B) Sequencing statistics for 2858 quality-filtered single cells. Black lines represent mean across all 2858 nuclei. For total reads, six outlier nuclei with very high read counts were omitted from the graph.

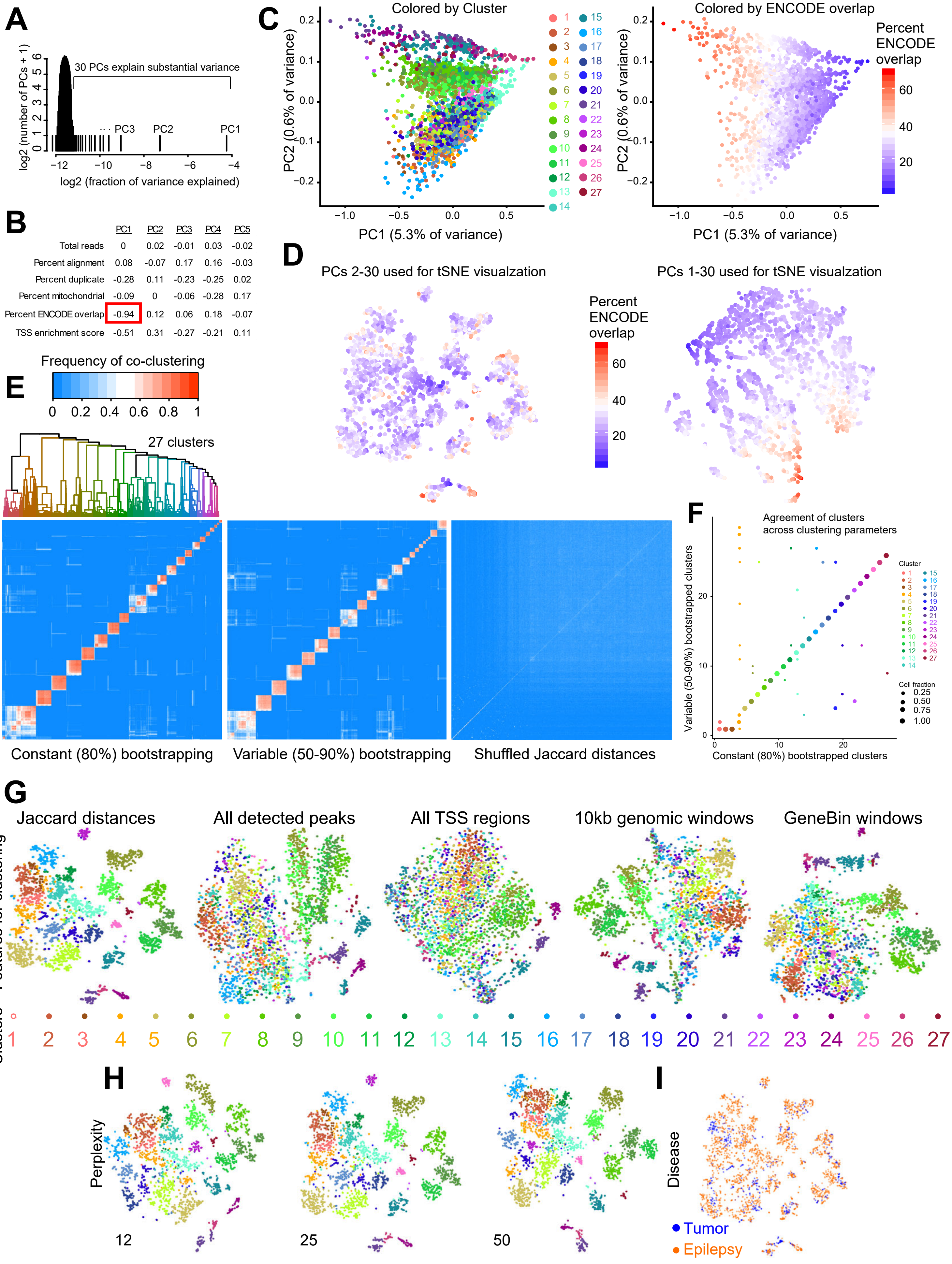

A
30 PCs explain substantial variance
log2 (number of PCs + 1)
log2 (fraction of variance explained)
PC3
PC2
PC1
B
PC1 PC2 PC3 PC4 PC5
Total reads 0 0.02 -0.01 0.03 -0.02
Percent alignment 0.08 -0.07 0.17 0.16 -0.03
Percent duplicate -0.28 0.11 -0.23 -0.25 0.02
Percent mitochondrial -0.09 0 -0.06 -0.28 0.17
Percent ENCODE overlap -0.94 0.12 0.06 0.18 -0.07
TSS enrichment score -0.51 0.31 -0.27 -0.21 0.11
C
Colored by Cluster
Colored by ENCODE overlap
PC1 (5.3% of variance)
PC2 (0.6% of variance)
Percent ENCODE overlap
D
PCs 2-30 used for tSNE visualzation
PCs 1-30 used for tSNE visualzation
Percent ENCODE overlap
E
Frequency of co-clustering
27 clusters
Bootstrapped clustering
Constant (80%) bootstrapping
Variable (50-90%) bootstrapping
Shuffled Jaccard distances
F
Agreement of clusters across clustering parameters
Variable (50-90%) bootstrapped clusters
Constant (80%) bootstrapped clusters
Cluster
Cell fraction
G
Jaccard distances
All detected peaks
All TSS regions
10kb genomic windows
GeneBin windows
Features for clustering
Clusters
H
Perplexity
12
25
50
I
Disease
Tumor
Epilepsy

**Supplementary Figure 4: High confidence clustering for single cell ATAC-seq data.**

A) Histogram showing the percentages of variance explained by each principal component of the Jaccard single cell distance matrix. The first 30 principal components explain substantial variance within the dataset.

B) Correlation of the first five principal components with quality metrics. Principal component 1 was omitted from further analysis due to strong negative correlation with ENCODE overlap.

C) Single nuclei evaluated by principal component analysis, with nuclei colored by cluster membership (*left*). Three major groups of nuclei were separated by PC2. Single nuclei were also colored by ENCODE overlap percentage, which is strongly negatively correlated with PC1 (*right*).

D) tSNE plot to visualize either principal components 2 to 30 (*left*) or 1 to 30 (*right*). Note, PCs 2 to 30 permit clear groupings with no ENCODE overlap gradient, whereas PCs 1 to 30 result in blurred cluster separations with a gradient of ENCODE spanning the clusters.

E) Bootstrapped iterative clustering to identify reproducible nuclear clusters. From the 2858 x 29 matrix of nuclei x principal component scores, we subsampled to either a constant 80% of nuclei (*left*) or a variable 50-90% of nuclei (*middle*), and calculated clusters using Jaccard-Louvain clustering (Tasic et al., 2018), which was repeated 200 times. Shuffled Jaccard distance matrix as input to PCA is shown (*right)*. Heatmaps display the frequency of co-clustering among nuclei. The constant 80% bootstrapping co-clustering matrix was used as input into Euclidean distance clustering, which yielded the final 27 clusters by cutting the tree to the major blocks of co-clustering nuclei. Nucleus order is not matched across the three plots.

F) Agreement between cluster memberships resulting from constant 80% bootstrapping and variable 50-90% bootstrapping, for most nuclei.

G) Visualization of nucleus groupings using five different feature sets (see Methods) using tSNE. Jaccard distances yielded clearest cell groupings. Cluster colors are applied in both (G) and (H).

H) Different perplexity parameters for tSNE visualization of cell groupings. Nuclear cluster groupings are evident at a wide range of perplexity values.

I) Visualization of disease status (tumor or epilepsy) for nuclei. Note that nuclei largely intermix regardless of disease status.

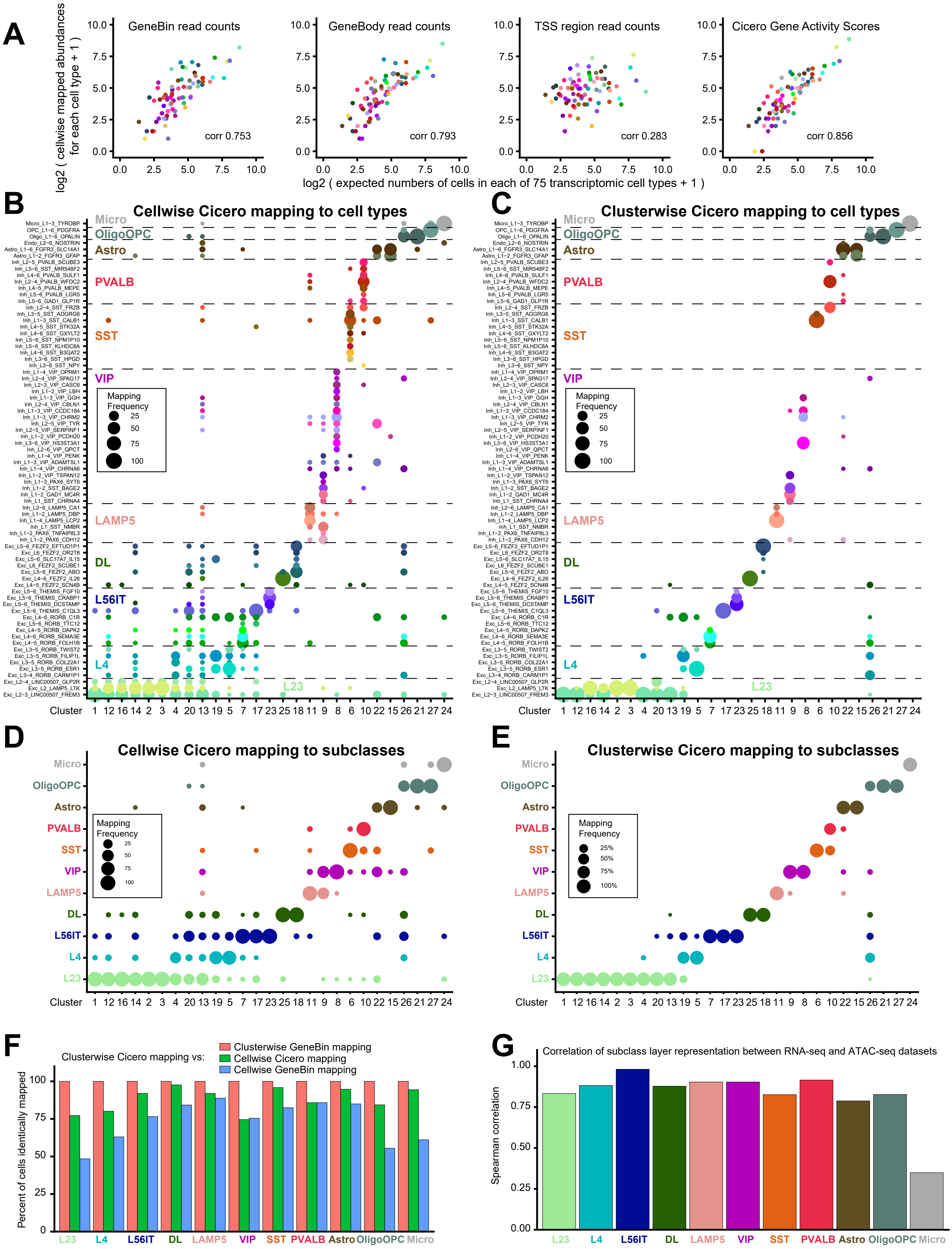
A
GeneBin read counts
corr 0.753
GeneBody read counts
corr 0.793
TSS region read counts
corr 0.283
Cicero Gene Activity Scores
corr 0.856
log2 ( cellwise mapped abundances for each cell type + 1 )
log2 ( expected numbers of cells in each of 75 transcriptomic cell types + 1 )
B
Cellwise Cicero mapping to cell types
C
Clusterwise Cicero mapping to cell types
Micro
OligoOPC
Astro
PVALB
SST
VIP
LAMP5
DL
L56IT
L4
L23
Mapping Frequency
25
50
75
100
Cluster 1 12 16 14 2 3 4 20 13 19 5 7 17 23 25 18 11 9 8 6 10 22 15 26 21 27 24
D
Cellwise Cicero mapping to subclasses
E
Clusterwise Cicero mapping to subclasses
25%
50%
75%
100%
F
Clusterwise Cicero mapping vs:
Clusterwise GeneBin mapping
Cellwise Cicero mapping
Cellwise GeneBin mapping
Percent of cells identically mapped
G
Correlation of subclass layer representation between RNA-seq and ATAC-seq datasets
Spearman correlation
L23 L4 L56IT DL LAMP5 VIP SST PVALB Astro OligoOPC Micro

**Supplementary Figure 5: Mapping ATAC-seq clusters to RNA-seq cell types.**

A) Expected abundances of each of the 75 transcriptomic cell types (Hodge et al., 2019), correlated with observed abundances of those cell types, using four different methods for computing gene-level information from each nucleus: *far left*: read counts in gene bins, *middle left*: read counts in gene bodies, *middle right:* read counts in 10kb-extended TSS regions, and *far right:* Cicero gene activity scores. Correlation values are Pearson correlation statistics between log-transformed expected and observed abundances plus one, for each of the 75 transcriptomic cell types. Computing gene-level information using cicero gene activity scores results in the greatest correlation between expectation and observation for cell type abundances.

B) Bootstrapped mapping of single nuclei (“cellwise”) to 75 transcriptomic cell types. Dot sizes indicate the frequencies of cell type mappings within each of the 27 ATAC-seq clusters.

C) Bootstrapped mapping of clusters (“clusterwise”) to 75 transcriptomic cell types. Dot sizes indicate the frequency each cluster maps to each transcriptomic cell type.

D) Bootstrapped mapping of single nuclei (“cellwise”) to 11 transcriptomic cell type subclasses. Dot sizes indicate the frequencies of subclass mappings within each of the 27 ATAC-seq clusters.

E) Bootstrapped mapping of clusters (“clusterwise”) to 11 transcriptomic cell type subclasses. Dot sizes indicate the frequency each cluster maps to each subclass. This plot represents the final mapped subclass assigned as the most frequent mapping for each cluster, which are used throughout the text.

F) Correlation of subclass mappings for all cells using four different mapping techniques. Overall, most cells are identically mapped to the same subclass with most of the techniques, with especially good agreement between both clusterwise mapping techniques.

G) Correlation between RNA-seq and ATAC-seq dataset layerwise distributions for the 11 subclasses. Most of the subclasses are observed in similar layer distributions in both datasets.

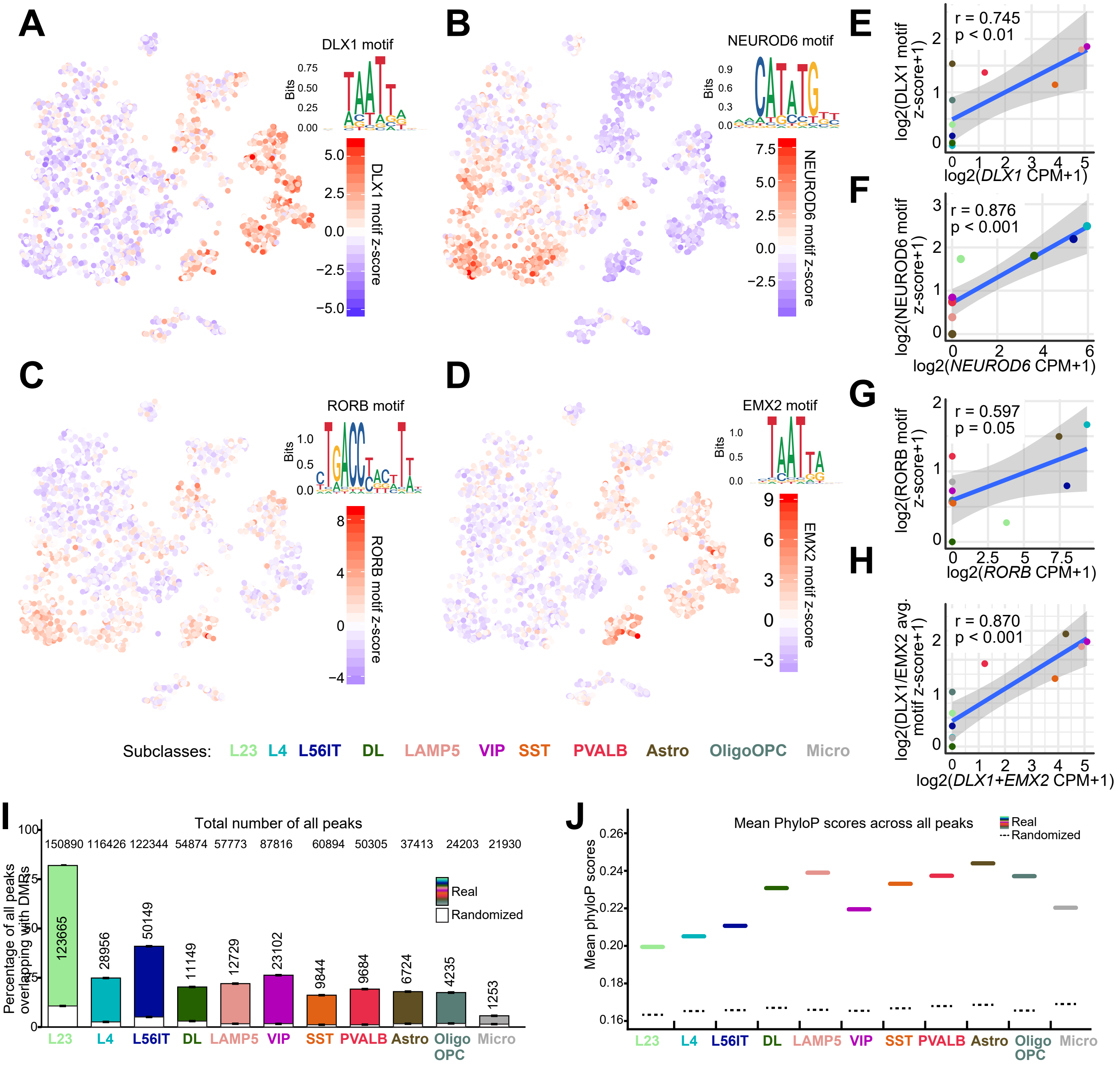

A
DLX1 motif
DLX1 motif z-score
B
NEUROD6 motif
NEUROD6 motif z-score
C
RORB motif
RORB motif z-score
D
EMX2 motif
EMX2 motif z-score
E
r = 0.745
p < 0.01
log2(DLX1 motif z-score+1)
log2(DLX1 CPM+1)
F
r = 0.876
p < 0.001
log2(NEUROD6 motif z-score+1)
log2(NEUROD6 CPM+1)
G
r = 0.597
p = 0.05
log2(RORB motif z-score+1)
log2(RORB CPM+1)
H
r = 0.870
p < 0.001
log2(DLX1/EMX2 avg. motif z-score+1)
log2(DLX1+EMX2 CPM+1)
Subclasses: L23 L4 L56IT DL LAMP5 VIP SST PVALB Astro OligoOPC Micro
I
Total number of all peaks
150890 116426 122344 54874 57773 87816 60894 50305 37413 24203 21930
Percentage of all peaks overlapping with DMRs
123665 28956 50149 11149 12729 23102 9844 9684 6724 4235 1253
Real
Randomized
L23 L4 L56IT DL LAMP5 VIP SST PVALB Astro Oligo OPC Micro
J
Mean PhyloP scores across all peaks
Real
Randomized
Mean phyloP scores
L23 L4 L56IT DL LAMP5 VIP SST PVALB Astro Oligo OPC Micro

**Supplementary Figure 6: Properties of human neocortical cell subclass-specific accessible genomic elements.**

A-D) Nuclei visualized by tSNE and colored by motif accessibilities for A) DLX1, B) NEUROD6, C) RORB, and D) EMX2 as calculated by chromVAR (Schep et al., 2017). Notice that DLX1 and EMX2 have similar recognition motifs and motif accessibility patterns among ATAC-seq cells. *DLX1* transcripts are specifically detected in inhibitory neurons and *EMX2* transcripts are specifically detected in astrocytes (Hodge et al., 2019).

E-H) Correlation between motif accessibilities and transcript abundances across cell subclasses for E) DLX1, F) NEUROD6, G) RORB, and H) the group of DLX1 and EMX2 (grouping by average for motif accessibility, and by sum for transcript abundances). Notice that EMX2 motif accessibility in astrocytes likely explains the spurious astrocytic DLX1 motif accessibility in absence of astrocytic DLX1 expression (Fig. S6E), resulting in improved ATAC-RNA correlation for the combination of DLX1 and EMX2, as versus DLX1 alone. r, pearson correlation coefficient. Two-tailed paired t-tests for significant correlation: DLX1 $t = 3.0$ df = 9 $p < 0.01$; NEUROD6 $t = 5.4$ df = 9 $p < 0.001$; RORB $t = 2.2$ df = 9 $p = 0.05$; DLX1 and EMX2 $t = 5.3$ df = 9 $p < 0.001$.

I) Percent overlap of ATAC-seq peaks with previously identified DMRs (Lister et al., 2013; Luo et al., 2017), comparing real peaks to randomized peak positions. Absolute numbers of detected peaks and peak-DMR overlaps are shown.

J) Mean phyloP scores across all peaks for cell subclass ATAC-seq peaks, compared to randomized peak positions.

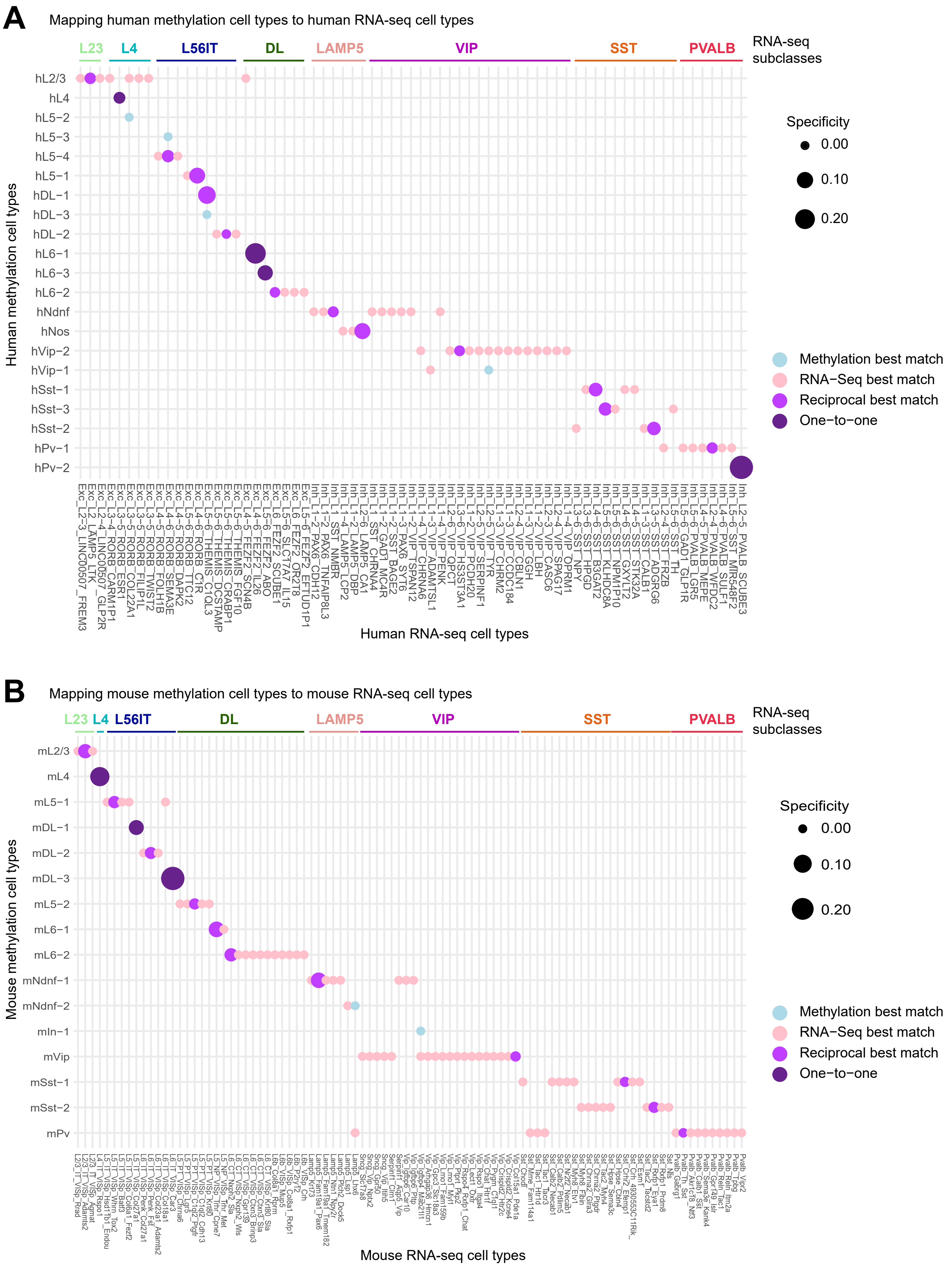

A
Mapping human methylation cell types to human RNA-seq cell types
L23
L4
L56IT
DL
LAMP5
VIP
SST
PVALB
RNA-seq subclasses
hL2/3
hL4
hL5−2
hL5−3
hL5−4
hL5−1
hDL−1
hDL−3
hDL−2
hL6−1
hL6−3
hL6−2
hNdnf
hNos
hVip−2
hVip−1
hSst−1
hSst−3
hSst−2
hPv−1
hPv−2
Human methylation cell types
Human RNA-seq cell types
Specificity
0.00
0.10
0.20
Methylation best match
RNA−Seq best match
Reciprocal best match
One−to−one
B
Mapping mouse methylation cell types to mouse RNA-seq cell types
L23
L4
L56IT
DL
LAMP5
VIP
SST
PVALB
RNA-seq subclasses
mL2/3
mL4
mL5−1
mDL−1
mDL−2
mDL−3
mL5−2
mL6−1
mL6−2
mNdnf−1
mNdnf−2
mIn−1
mVip
mSst−1
mSst−2
mPv
Mouse methylation cell types
Mouse RNA-seq cell types
Specificity
0.00
0.10
0.20
Methylation best match
RNA−Seq best match
Reciprocal best match
One−to−one

**Supplementary Figure 7: Mapping methylation cell types to RNA-seq cell types.**

Mouse and human cell types derived from mCH methylation data (Luo et al., 2017), mapped to RNA-seq derived cell types from human MTG (A) (Hodge et al., 2019) or mouse V1 (B) (Tasic et al., 2018). Specificity is defined as the difference between the best correlation and the second-best correlation among all the competing choices. For analysis in Figs S6 and S8, the methylation cell types were aggregated into transcriptomically defined subclasses according to their best matches shown here.

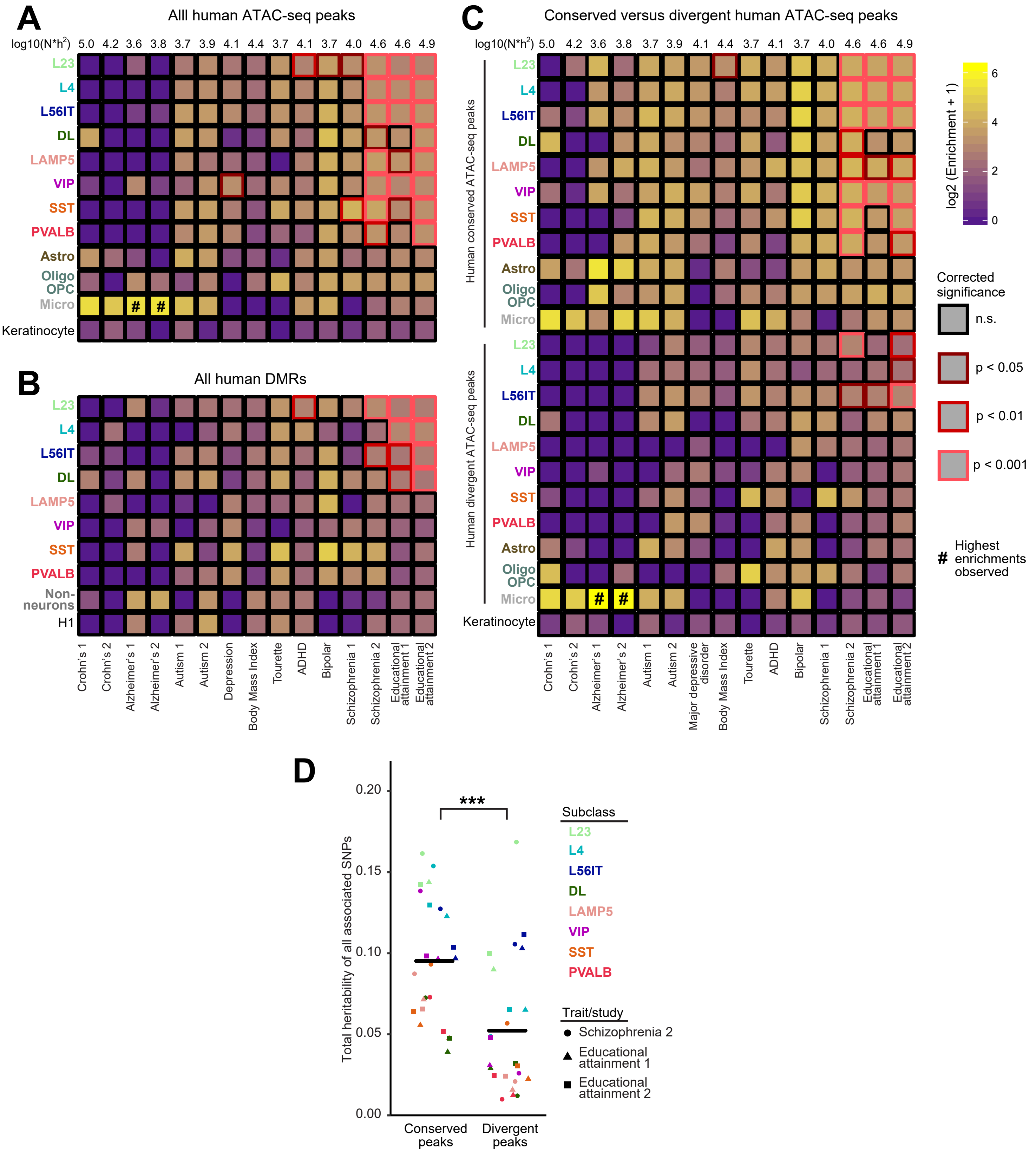
A
Alll human ATAC-seq peaks
log10(N*h2)
5.0 4.2 3.6 3.8 3.7 3.9 4.1 4.4 3.7 4.1 3.7 4.0 4.6 4.6 4.9
L23
L4
L56IT
DL
LAMP5
VIP
SST
PVALB
Astro
Oligo OPC
Micro
Keratinocyte
#
B
All human DMRs
L23
L4
L56IT
DL
LAMP5
VIP
SST
PVALB
Non-neurons
H1
Crohn's 1
Crohn's 2
Alzheimer's 1
Alzheimer's 2
Autism 1
Autism 2
Depression
Body Mass Index
Tourette
ADHD
Bipolar
Schizophrenia 1
Schizophrenia 2
Educational attainment 1
Educational attainment 2
C
Conserved versus divergent human ATAC-seq peaks
log10(N*h2)
5.0 4.2 3.6 3.8 3.7 3.9 4.1 4.4 3.7 4.1 3.7 4.0 4.6 4.6 4.9
Human conserved ATAC-seq peaks
Human divergent ATAC-seq peaks
L23
L4
L56IT
DL
LAMP5
VIP
SST
PVALB
Astro
Oligo OPC
Micro
Keratinocyte
Crohn's 1
Crohn's 2
Alzheimer's 1
Alzheimer's 2
Autism 1
Autism 2
Major depressive disorder
Body Mass Index
Tourette
ADHD
Bipolar
Schizophrenia 1
Schizophrenia 2
Educational attainment 1
Educational attainment 2
log2 (Enrichment + 1)
0
2
4
6
Corrected significance
n.s.
p < 0.05
p < 0.01
p < 0.001
### Highest enrichments observed
D
Total heritability of all associated SNPs
0.00
0.05
0.10
0.15
0.20
***
Conserved peaks
Divergent peaks
Subclass
L23
L4
L56IT
DL
LAMP5
VIP
SST
PVALB
Trait/study
Schizophrenia 2
Educational attainment 1
Educational attainment 2

**Supplementary Figure 8: ATAC-seq peaks uncover strong associations between diseases and conserved accessible elements for human neocortical neuron subclasses.**

A-B) Associations between genome-wide association study diseases/traits and subclass ATAC-seq peaksets (A) and methylation DMRs (B, Lister et al., 2013; Luo et al., 2017). Heatmap fill colors represent enrichments, which are defined as the ratio of the proportion of heritability contained by that peakset's linked SNPs, to the proportion of that peakset's linked SNPs, as calculated by LDSC (Bulik-Sullivan et al., 2015; Finucane et al., 2015). Red outlines demonstrate significant associations after performing Bonferroni correction for multiple hypothesis testing (180 tests for ATAC-seq peaks and 150 tests for DMRs). Widespread associations between multiple brain diseases and multiple cortical neuron subclasses are observed with both ATAC-seq peaks and DMRs. Strongest associations are seen for Alzheimer's disease in microglial ATAC-seq peaks, and significant associations are seen for educational attainment and schizophrenia across multiple neuronal classes.

C) Associations between conserved (also observed in mouse ATAC-seq) and divergent (not observed in mouse ATAC-seq) human ATAC-seq peaks, and genome-wide association study diseases/traits. Conserved human peaks display generally greater enrichment and more significant associations than do divergent human peaks, in particular for educational attainment and schizophrenia. A notable exception is microglia in Alzheimer's disease, which shows greater enrichment in divergent peaks, although this enrichment does not pass statistical significance, possibly due to low overall total heritability and hence statistical power in Alzheimer's studies. Bonferroni-corrected p-values are employed (345 tests performed).

D) Total summed heritability of all SNPs associated with conserved peaks, versus those associated with divergent peaks, for three studies with multiple significant neuron subclass associations. ***$p < 0.01$ by heteroscedastic t-test, $t = 3.8$, $df = 45.6$.

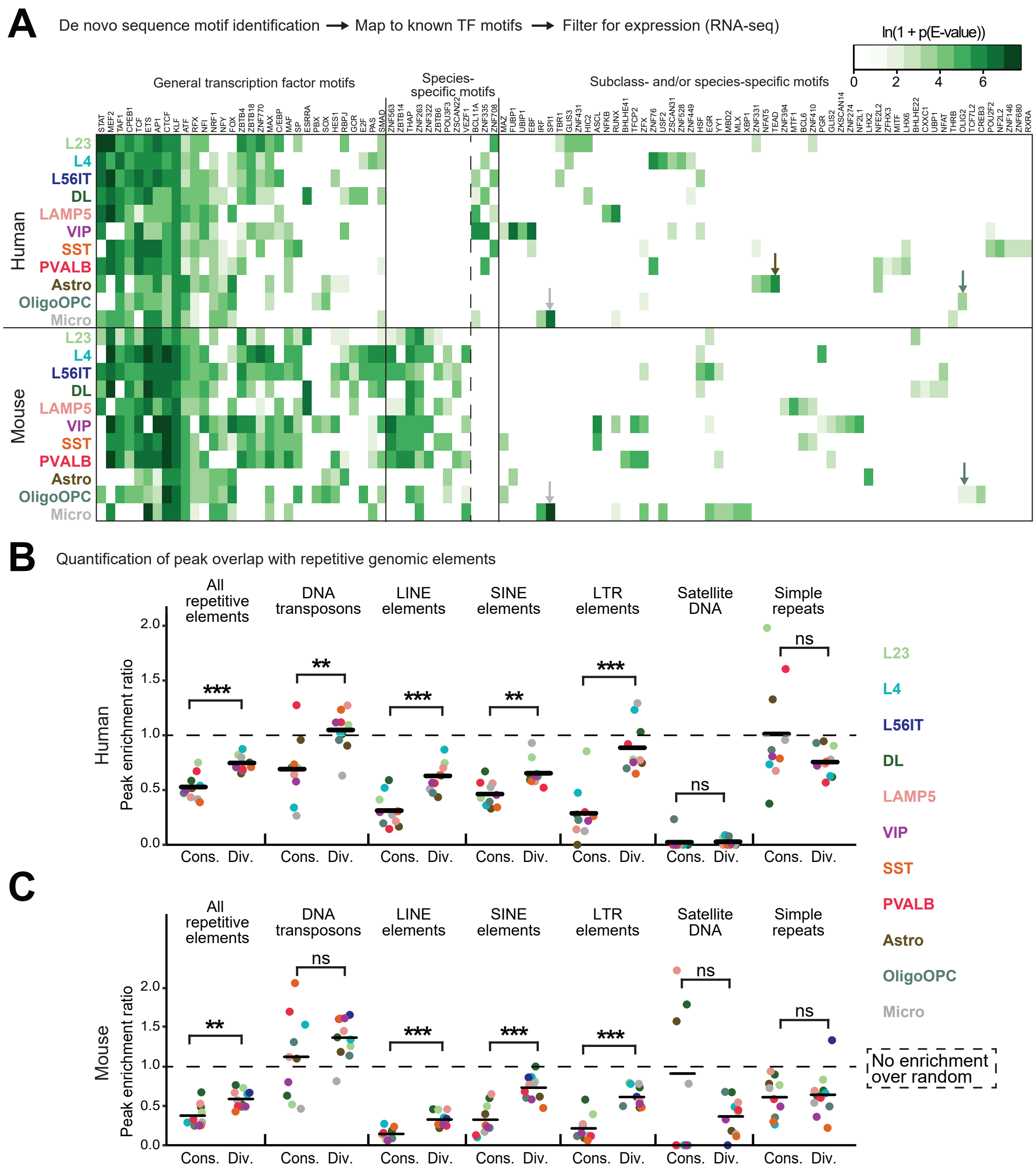
A
De novo sequence motif identification → Map to known TF motifs → Filter for expression (RNA-seq)
ln(1 + p(E-value))
0
2
4
6
General transcription factor motifs
Species-specific motifs
Subclass- and/or species-specific motifs
Human
Mouse
L23
L4
L56IT
DL
LAMP5
VIP
SST
PVALB
Astro
OligoOPC
Micro
B
Quantification of peak overlap with repetitive genomic elements
All repetitive elements
DNA transposons
LINE elements
SINE elements
LTR elements
Satellite DNA
Simple repeats
Peak enrichment ratio
Cons.
Div.
***
**
ns
C
No enrichment over random

**Supplementary Figure 9: Accessible genomic elements disclose active chromatin regulators and mechanisms for neocortical gene expression changes across mammalian species.**

A) Active transcriptional regulators in human and mouse brain cell subclasses, revealed by motifs in ATAC-seq peaks and gene expression by transcriptomics (Tasic et al., 2018; Hodge et al., 2019). E-value indicates the p-value from Fisher's exact test, corrected for multiple testing as calculated by MEME-CHIP. Arrows indicate strong and specific microglial enrichments for SPI1/PU.1 (gray) and for TEAD in human astrocytes (brown) and for OLIG2 in oligodendrocytes/OPCs (cadet blue).

B-C) Overlap of conserved or divergent peaks by cell subclass with multiple classes of repetitive genomic elements in both human (B) and mouse (C) sn/scATACseq datasets. An enrichment value of 1.0 corresponds to no fold change between real and random peak enrichment. Black bars represent the mean across the eleven neocortical cell subclasses. Heteroscedastic t-tests: *** $p < 0.001$, ** $p < 0.01$, ns not significant. Human all elements $t = 5.2$, df = 14.5; human DNA transposons $t = 3.4$, df = 14.8; human LINE $t = 5.3$, df = 18.3; human SINE $t = 3.8$, df = 18.8; human LTR $t = 6.1$, df = 18.3; human satellite $t = 0.2$, df = 12.4; human simple $t = 1.6$, df = 10.1; mouse all elements $t = 3.7$, df = 16.9; mouse DNA transposons $t = 1.3$, df = 12.7; mouse LINE $t = 5.2$, df = 18.5; mouse SINE $t = 5.2$, df = 18.0; mouse LTR $t = 6.1$, df = 17.5; mouse satellite $t = 1.6$, df = 9.7; mouse simple $t = 0.3$, df = 19.0.

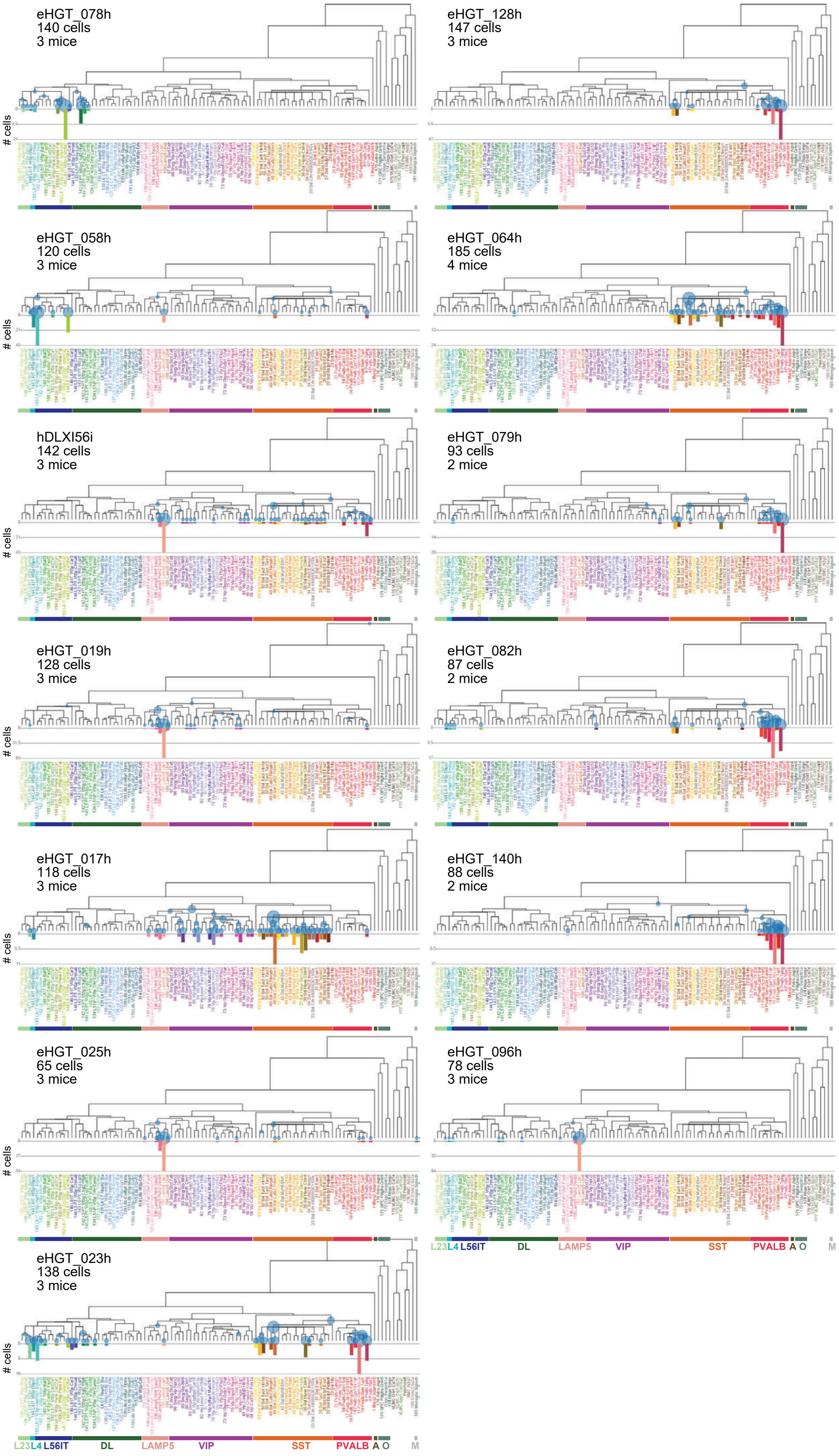
eHGT_078h
140 cells
3 mice
eHGT_128h
147 cells
3 mice
eHGT_058h
120 cells
3 mice
eHGT_064h
185 cells
4 mice
hDLXI56i
142 cells
3 mice
eHGT_079h
93 cells
2 mice
eHGT_019h
128 cells
3 mice
eHGT_082h
87 cells
2 mice
eHGT_017h
118 cells
3 mice
eHGT_140h
88 cells
2 mice
eHGT_025h
65 cells
3 mice
eHGT_096h
78 cells
3 mice
eHGT_023h
138 cells
3 mice
### cells
L23 L4 L56IT
DL
LAMP5
VIP
SST
PVALB A O
M

**Supplementary Figure 10: Cell type validation of enhancer-AAV-labeled cells via scRNA-seq.** Numbers of sorted labeled cells with each enhancer-AAV vector shown in Figures 3 and 4 and S11, mapped to the cell type transcriptomic taxonomy of mouse V1 (Tasic et al., 2018). Dendrogram leaves represent 111 transcriptomic cell types. Circles on the dendrogram represent the number of cells that could be mapped to that point in the dendrogram (starting from the root) and bar plots below the leaves represent the number of each cell type recovered that mapped to that final leaf.

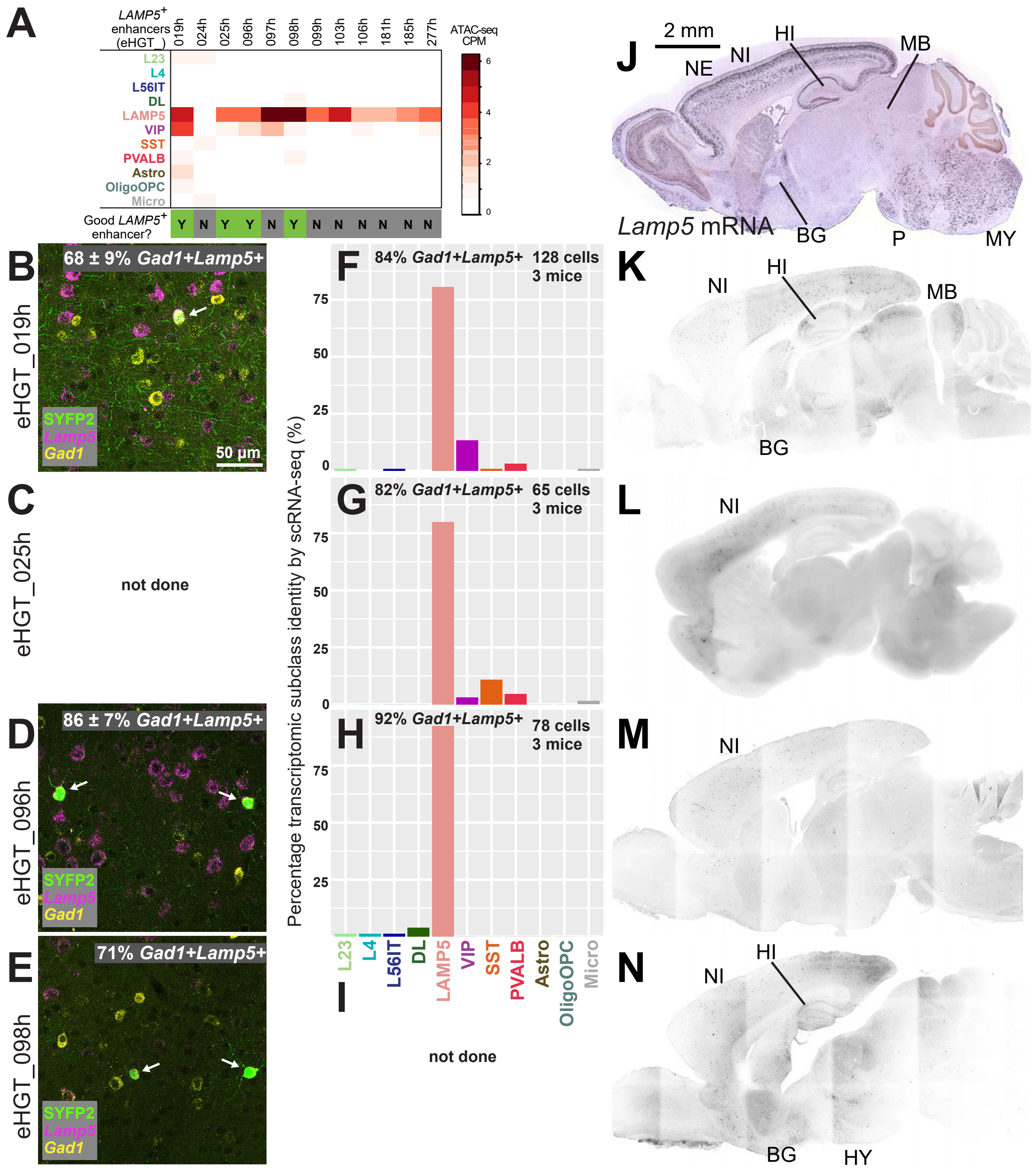
A
LAMP5+ enhancers (eHGT_)
019h 024h 025h 096h 097h 098h 099h 103h 106h 181h 185h 277h
ATAC-seq CPM
L23
L4
L56IT
DL
LAMP5
VIP
SST
PVALB
Astro
OligoOPC
Micro
Good LAMP5+ enhancer?
Y N Y Y N Y N N N N N N
B
eHGT_019h
68 ± 9% Gad1+Lamp5+
SYFP2
Lamp5
Gad1
50 µm
C
eHGT_025h
not done
D
eHGT_096h
86 ± 7% Gad1+Lamp5+
E
eHGT_098h
71% Gad1+Lamp5+
F
84% Gad1+Lamp5+ 128 cells 3 mice
G
82% Gad1+Lamp5+ 65 cells 3 mice
H
92% Gad1+Lamp5+ 78 cells 3 mice
Percentage transcriptomic subclass identity by scRNA-seq (%)
I
not done
J
2 mm
Lamp5 mRNA
NE
NI
HI
MB
BG
P
MY
K
L
M
N
HY

**Supplementary Figure 11: A collection of LAMP5 subclass-specific enhancer-AAV vectors.**

A) Twelve putative LAMP5 enhancers identified from snATAC-seq data and cloned into AAV vectors. Four of the twelve (33%) exhibited high selectivity for LAMP5 cells in mouse retro-orbital assay (indicated with green boxes).

B-E) Positive labeling of $Gad1^+Lamp5^+$ cells (arrows) by each enhancer-AAV vector, as demonstrated by mFISH HCR in V1 L2/3. Percentages indicate the specificity of SYFP2 labeling for $Gad1^+Lamp5^+$ cells (n = 3 mice for eHGT_019h and 096h, and n = 1 for eHGT_098h).

F-I) scRNA-seq in V1 confirms the transcriptomic cell subclass identity of enhancer-AAV vector-labeled cells. Bargraph shows the percentage of single cells that map to a transcriptomic cell type within that subclass. In contrast, the percentages given in text are the percentage of cells recovered that expressed *Gad1* and *Lamp5*.

J) *Lamp5* mRNA expression pattern (Allen Institute public ISH data) shows multiple sites of expression throughout mouse brain. Abbreviations: NE neocortical excitatory neurons, NI neocortical inhibitory neurons, HI hippocampal Inh neurons, BG basal ganglia, MB midbrain, MY medulla, P pons, HY hypothalamus.

K-N) LAMP5-selective enhancers eHGT_025h and 096h label only *Gad1+Lamp5+* cells in neocortex, but eHGT_019h and 098h also label various subcortical brain regions also seen in the endogenous *Lamp5* mRNA expression pattern. Confocal images are shown except for L which is an epifluorescence image.
